# Apolipoprotein L9 interacts with LC3/GABARAP and is a microtubule-associated protein with a widespread subcellular distribution

**DOI:** 10.1101/671065

**Authors:** Arvind A. Thekkinghat, Kamlesh K. Yadav, Pundi N. Rangarajan

## Abstract

Mouse Apolipoprotein L9 is a 34-kDa phosphatidylethanolamine (PE)-binding protein. The gene is present only in mouse and rat genomes; hence it is taxonomically restricted. To understand why, it is essential to uncover details about its functions in cellular processes. Here we show that ApoL9 interacts with the proteins of the LC3 and GABARAP subfamilies, which are key players in macroautophagy. In amino-acid starved cells it preferentially interacts with lipidated LC3B, likely by binding to its PE moiety. On treatment with autophagy inhibitors bafilomycin A1 and chloroquine, ApoL9 is found near swollen mitochondria and on lysosomes/LAMP1-positive compartments. However, ApoL9 itself does not seem to be degraded as a result of autophagy, suggesting that it is not an autophagy cargo receptor. Deletions in a putative transmembrane region between amino acids 110 and 145 abolish PE-binding. In addition, ApoL9 can redistribute to stress granules, can homooligomerize, and is a microtubule-associated protein. In short, its distribution in the cell is quite widespread, suggesting that it could have functions at the intersection of membrane binding and reorganization, autophagy, cellular stress and intracellular lipid transport.

**Summary statement:** This article is about how Apolipoprotein L9, a lipid-binding protein, has versatile properties and influences a variety of processes taking place inside an animal cell.

## Introduction

Mouse ApoL9, encoded by two independent genes *Apol9a* and *Apol9b* on chromosome 15, has previously been shown to have either antiviral or pro-viral effects during infection of cells by different types of viruses (Kreit et al., 2014; Kreit et al., 2015; Arvind and Rangarajan, 2016). Expression of ApoL proteins is also induced by interferons and TNF-α (Zhaorigetu et al., 2011, Monajemi et al., 2002). Small quantities of ApoL9 secreted from macrophages during interferon induction have been shown to promote epithelial cell proliferation in a paracrine fashion (Sun et al., 2015). However, most of the protein is retained within the cell. In a previous study, we used B16F10 melanoma cells to look at the basic expression pattern of constructs expressing ApoL9, examined its levels in various mouse tissues, and viewed it in the context of infection by Japanese Encephalitis virus (Arvind and Rangarajan, 2016). We reported that ApoL9 is a phosphatidylethanolamine-binding protein that, in normal conditions, has a general cytoplasmic distribution and can localize to ubiquitin-positive bodies called ALIS (aggresome-like induced structures) and aggresomes. ApoL9 is expressed at moderate to high levels in mouse liver and brain, suggesting some function of relevance for the protein in these major tissues.

In order to understand the functions of ApoL9, it is essential to know more about its distribution in the cell and the proteins it interacts with. In this study, our objective is to attempt to uncover as many clues as possible to help place ApoL9 in the context of processes taking place in the cell. Since phosphatidylethanolamine has a unique function as a modifier of autophagosome marker proteins Atg8 and its orthologues (Kabeya 2004), we examined whether ApoL9 could influence autophagy. We investigate how ApoL9 interacts with PE by screening deletion mutants of the protein for PE-binding. We also investigate the fate of ApoL9 when cells are subjected to stress, by subjecting cells to treatments that induce cell stress, and observe the distribution and levels of ApoL9 under these conditions. We find that ApoL9 is a dynamic protein that localizes to various compartments in the cell under different conditions.

## Results

### 1. ApoL9 interacts with the mammalian orthologues of Atg8

We previously reported that ApoL9 localizes to ALIS-like structures, which also contain LC3 and SQSTM1 (sequestosome 1/p62), proteins that have key roles in autophagic processes (Arvind and Rangarajan, 2016). ApoL9 also binds PE, whose covalent modification of LC3 is a crucial event in the initiation and progression of macroautophagy (henceforth referred to as autophagy). Several proteins that regulate autophagy interact with LC3 and its homologues, which are central players in autophagy (Wild et al., 2013). We investigated to check if ApoL9 could also interact with any of these proteins. Mouse *Lc3a, Lc3b, Gabarap, Gabarapl1*, and *Gabarapl2* were expressed as Glutathione S-transferase fusion proteins in *E.coli*, bound to glutathione-agarose, and examined for their ability to interact with V5-tagged ApoL9 (ApoL9-V5) expressed in HEK293T cells. SDS-PAGE followed by immunoblotting with anti-V5 antibody revealed that ApoL9 interacts with all the members of the LC3 and GABARAP subfamilies, and that the interaction was strongest with GABARAP and GABARAPL1 (**Fig. 1A**). To validate this interaction in vivo, we attempted to co-immunoprecipitate GABARAP along with ApoL9. Using anti-V5 agarose, GABARAP could be immunoprecipitated along with ApoL9-V5 from HEK293Tcells transfected with both *Apol9-V5* and *GFP-Gabarap* constructs (**Fig. 1B**). GFP-LC3B could also be co-immunoprecipitated with ApoL9-V5 in a similar fashion, indicating that ApoL9 interacts with both these proteins inside the cell.

**Fig.1:**
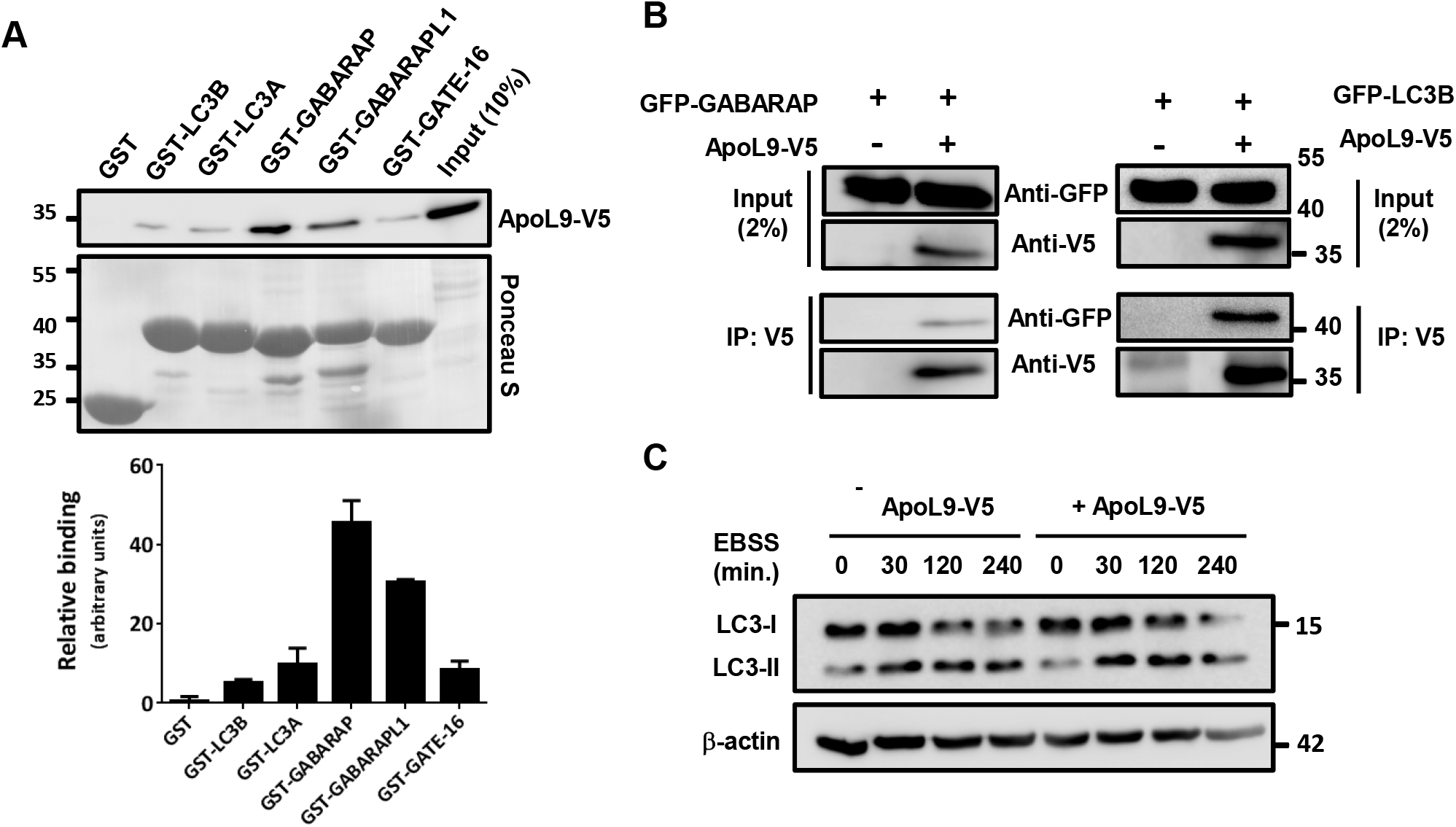
ApoL9 interacts with proteins of the LC3 and GABARAP subfamilies. **A.** Interaction of ApoL9-V5 expressed in HEK293T cells with recombinant GST-tagged proteins of the LC3 and GABARAP subfamilies. GST served as a negative control. GST-fusion proteins were stained by Ponceau S. Quantification of binding is shown in the graph below. Values represent mean ± s.d.; n=3. **B.** Co-immunoprecipitation of GFP-tagged LC3B and GABARAP with ApoL9-V5, from lysates of HEK293T cells transfected with *ApoL9-V5* and either of the GFP-fusions, by anti-V5 agarose affinity gel. Cells transfected with control vector and *GFP-Gabarap/GFP-Lc3b* served as negative controls. 2% of the total quantity of protein lysate taken for IP served as input to confirm efficient transfection. **C**. Comparison of LC3-I and LC3-II levels in HEK293T cells expressing (lanes 5-8) and not expressing (lanes 1-4) ApoL9-V5. *Apol9-V5* was transfected into cells by the calcium phosphate method. Cells transfected with empty vector (insert-less *pBApo-EF1α pur)* served as control. β-actin was used as a loading control.

Since the conversion of the cytosolic form of LC3 (LC3-I) to LC3-PE conjugate (LC3-II) occurs at the onset of autophagy (Kabeya, 2000), we investigated if overexpression of ApoL9 had any influence on levels of lipidated LC3 (LC3-II). HEK293T cells were transfected with either control plasmid or plasmid expressing *Apol9-V5*, and to induce autophagy the cells were starved of amino acids by culturing in Earle’s Balanced Salt Solution (EBSS) for different time periods. Immunoblotting did not reveal any significant difference in levels of LC3-II between the two conditions (**Fig. 1C**), suggesting that ApoL9 expression does not alter levels of lipidated LC3, at least under conditions of amino acid starvation.

### 2. ApoL9 localises to mitochondria and LAMP1-positive structures during treatment with compounds that inhibit autophagy/induce mitochondrial damage

When B16F10 melanoma cells stably expressing ApoL9-V5 (B16F10^L9^) cells were treated with bafilomycin A1, a V-ATPase inhibitor which inhibits the degradative capacity of lysosomes (Mauvezin et al., 2015), the distribution of ApoL9 in the cell was significantly altered. Starting approximately 8–10 h after treatment, three different patterns could be observed –localisation in the juxtanuclear region, ring-like structures in the cytoplasm, and thread-like, highly curved/entangled structures in the cytoplasm at later time points (**Fig. 2A**). The juxtanuclear ApoL9 co-localized with structures stained by an anti-LAMP1 antibody, indicating that they were lysosomes or other LAMP1-positive structures (**Fig. 2B**). The ring-like structures positive for ApoL9 increased in number with time, and co-localized with the outer mitochondrial marker protein TSPO, indicating that they were swollen mitochondria (**Fig. 2C**). The curved, thread-like structures also increased in number with time, but did not co-localize with any organelle marker we used (**Fig. 2B, C**). These three patterns are not mutually exclusive, but we often notice that cells in which juxtanuclear lysosomal localization predominates contain fewer curved, thread-like structures and vice versa. Thus, the possibility these two patterns are linked cannot be ruled out.

**Fig.2:**
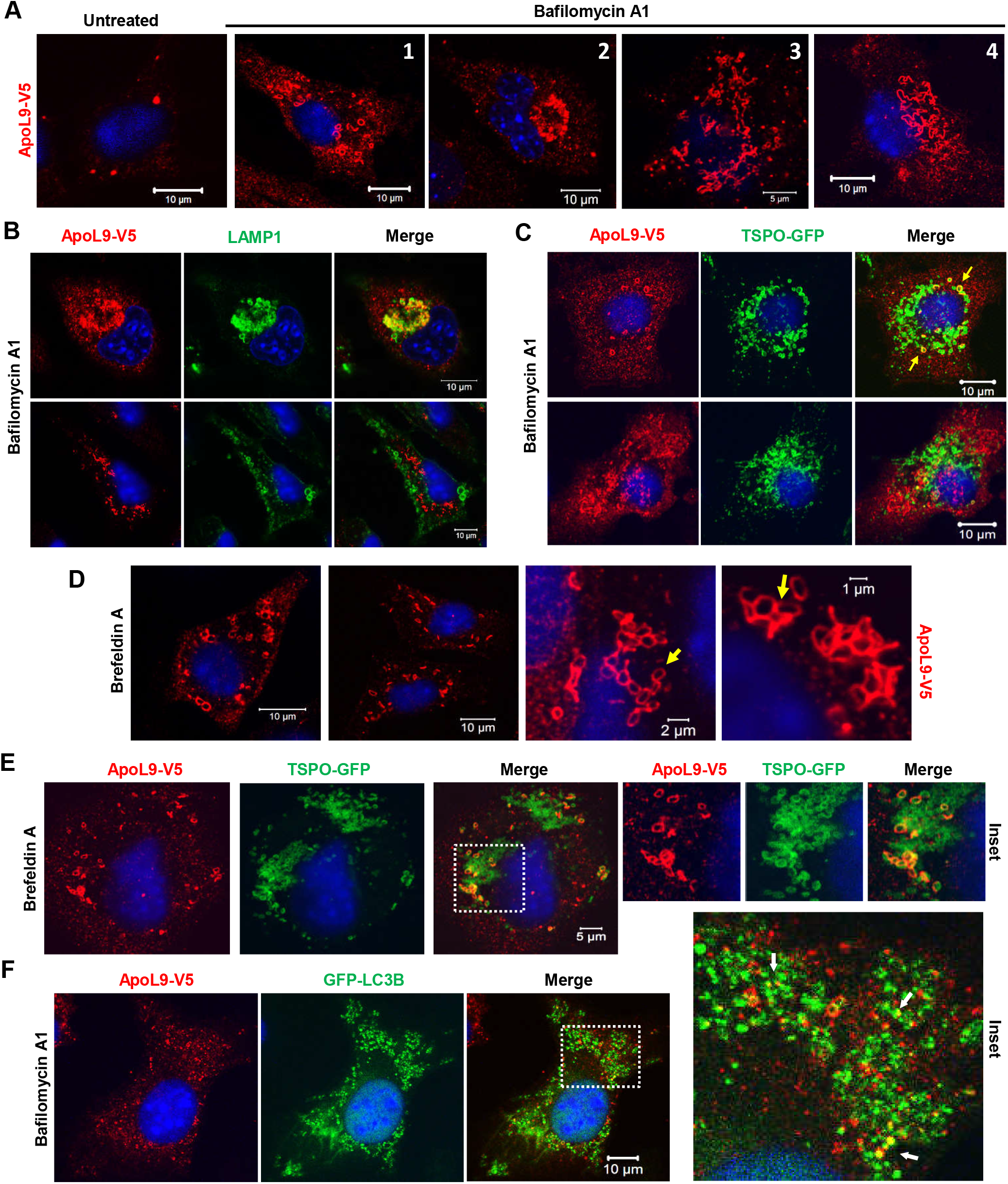
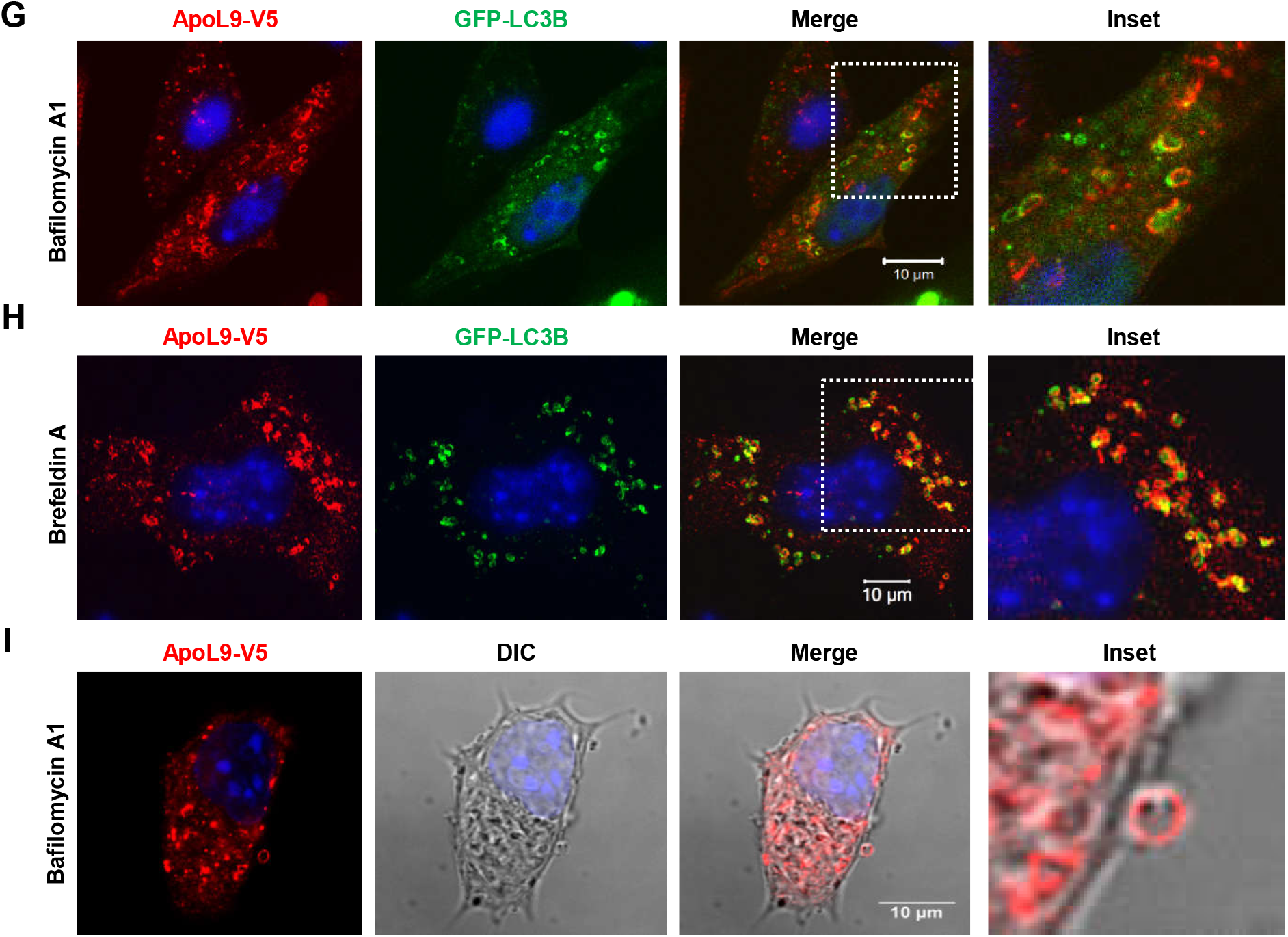
Subcellular distribution of ApoL9 after treatment with bafilomycin A1 and brefeldin A. **A.** Patterns of distribution of ApoL9-V5 in various cells of B16F10^L9^ after treatment with 200 nM bafilomycin A1 for 10-18 h, showing: ring-shaped structures (1), accumulation in the juxtanuclear region (2), and curved, thread-like structures (3, 4). Anti-V5 antibody was used to detect ApoL9. **B.** Indirect immunofluorescence for ApoL-V5 and lysosomal marker LAMP1 in B16F10^L9^ cells treated with bafilomycin A1 for 12 h, showing co-localization in the juxtanuclear region (upper panel). ApoL9 in curved, thread-like structures (lower panel) does not co-localize with LAMP1. Anti-V5 and anti-LAMP1 antibodies were used. **C.** Indirect immunofluorescence for ApoL9 and outer mitochondrial membrane (OMM) marker TSPO in B16F10^L9^cells treated with bafilomycin A1 for 12 h, showing co-localization of ApoL9-V5-positive rings with TSPO-GFP (upper panel, yellow arrows). ApoL9-V5 in curved, threadlike structures (lower panel) does not co-localize with TSPO-GFP. *Tspo-GFP* was electroporated into B16F10^L9^ cells. **D**. Patterns of distribution of ApoL9-V5 in various cells after treatment with 10 mg/mL brefeldin A for 14-18 h. Note the proximity between ring-shaped structures (yellow arrows) in images of higher magnification. **E.** Indirect immunofluorescence for ApoL9-V5 and TSPO-GFP in B16F10^L9^ cells treated with brefeldin A, and (right) magnified inset. Note that only a fraction of the mitochondria are positive for ApoL9. **F**. A B16F10^L9^ cell electroporated with the *GFP-Lc3b* construct and treated with bafilomycin A1 for 12 h, showing GFP-LC3B on punctate autophagosomes. ApoL9-V5 is not present on autophagosomes, but partial co-localization is visible in a few places (magnified boxed inset; white arrows). **G-H.** Images showing GFP-LC3B (transfected by electroporation) in the vicinity of ApoL9-containing mitochondria in B16F10^L9^ cells treated with bafilomycin A1 and brefeldin A, respectively. Boxed insets are magnified. **I.** An ApoL9-positive ring-shaped vesicle exiting the cell boundary. B16F10^L9^ cells were treated with bafilomycin A1 for 16h.

We also noticed ApoL9 localizing to ring-like mitochondria, starting approximately 8-10 h post treatment with brefeldin A (**Fig. 2D**), an ER stress inducer that inhibits protein transport from the ER to the Golgi apparatus by inhibiting a guanine nucleotide exchange factor GBF1, causing a collapse of the proteins into the ER (Fujiwara et al., 1988; Niu et al., 2005). ApoL9 was only found on some mitochondria, but not all (**Fig. 2E**), during treatment by either brefeldin A or bafilomycin A1, and this number steadily increased with incubation time with these compounds. ER stress induced by brefeldin A has previously been shown to damage mitochondria by depolarizing them, triggering apoptosis by the classical pathway (Lee et al., 2013). Bafilomycin A1 has been demonstrated to be a potassium ionophore that leads to impairment of mitochondrial functions by causing them to swell and lose membrane potential (Teplova et al., 2007). These properties of the compounds, and the considerable time period after which ApoL9 localises to mitochondria in B16F10^L9^ cells during treatment with these compounds, are indications that the ApoL9-positive mitochondria might have suffered damage to their integrity.

There are clear differences between ApoL9 localization during treatment with bafilomycin A1 and brefeldin A. During treatment with brefeldin A, ApoL9 localizes only to TSPO-positive ring-like structures which are swollen mitochondria. These are often seen clustered together, appearing to be in contact (**Fig. 2D**). No localization to lysosomes (**Fig. S1.A**) or to curved thread-like structures can be seen. Also, the fluorescence signal from ApoL9 becomes weaker with time, unlike in the case of bafilomycin A1 treatment.

In bafilomycin A1-treated cells, LC3B (as GFP-LC3B) accumulates in numerous punctate autophagosomes due to the downstream block in autophagy. ApoL9 does not localize to these structures, suggesting that it is not a part of the autophagosome per se, though transient interactions with autophagosomes might occur (**Fig. 2F**). Upon treatment with bafilomycin A1 and brefeldin A, LC3B and GABARAP also appear in the vicinity of mitochondria along with ApoL9, which might indicate that the autophagic machinery is recruited to the damaged mitochondria (**Fig.2G, H, and S1.B**). In some cases there appears to be only partial colocalization between ApoL9 and TSPO/LC3B, suggesting that they might be part of more than one structure on or near the mitochondria. But spatial restriction of LC3B/GABARAP localization to ApoL9-positive mitochondria is very clearly evident. The presence of ApoL9 on LAMP-1-positive structures and swollen mitochondria when autophagy is inhibited, and its co-localization with LC3B and GABARAP, suggests a role in autophagy.

Another phenomenon observed in cells treated with bafilomycin A1 is the presence of a number of the ApoL9-positive ring-like structures outside the boundaries of the cells that once contained them. They could be seen well outside the boundary of the cells, exiting the membrane (**Fig. 2I**), or in close contact with the membrane (before exit), in various images (**Fig. S1.C,D**). Mitochondria have previously been described to be ejected from cells undergoing phenomena such as necroptosis and serve as a source of DAMPs (damage-associated molecular patterns) (Maeda and Fadeel, 2014; Krysko et al., 2011). This study is the first report of ApoL9-containing mitochondria in the extracellular milieu, though it is not clear how much the mitochondria inside the cells and mitochondria outside the cells differ in membrane integrity and protein composition. We didn’t observe this phenomenon in cells treated with brefeldin A.

### 3. ApoL9 accumulates in the Triton X-100-insoluble fraction of lysates of cells treated with bafilomycin A1, but does not seem to be a receptor for selective autophagy

In western blots of cell lysates of B16F10^L9^ cells treated with bafilomycin A1, there is a gradual decrease of ApoL9 in the Triton X-100 soluble fraction and a corresponding increase in the insoluble fraction (**Fig. 3A**), which correlates with the gradual accumulation of ring-like, curved or thread-like entangled structures in the cytoplasm over the same time period.

**Fig.3:**
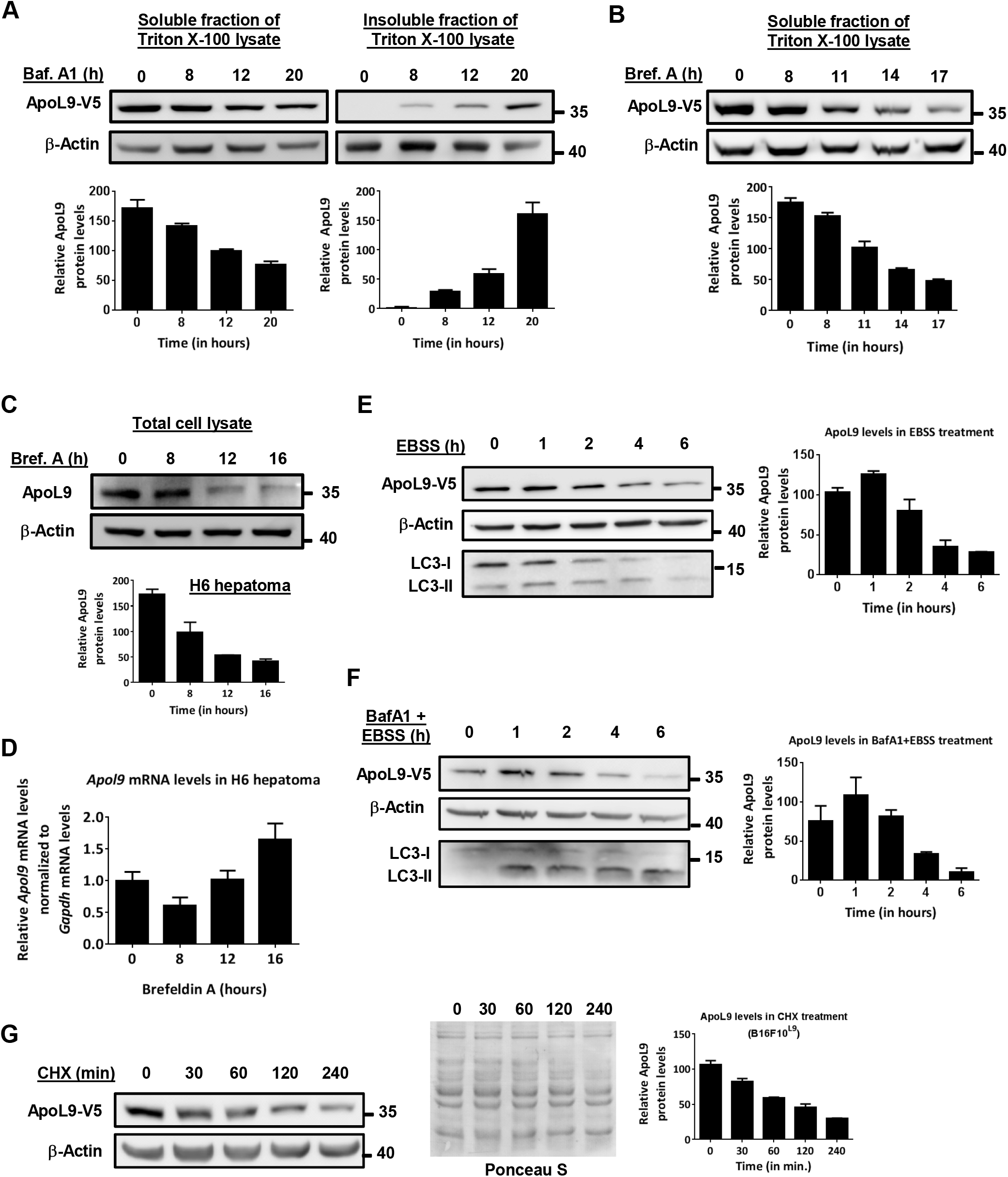
Effect of treatment with various compounds on ApoL9 protein levels in the cell. **A.** Immunoblotting for ApoL9-V5 in Triton X-100-soluble and insoluble fractions of B16F10^L9^cells treated with bafilomycin A1 for the indicated time periods, and (below) quantification of the blots. **B.** Immunoblotting for ApoL9-V5 in the Triton X-100-soluble fraction of B16F10^L9^ cells treated with brefeldin A for the indicated time periods and (below) quantification. No ApoL9 was detected in the Triton X-100-insoluble fraction. **C.** Immunoblotting for endogenous ApoL9 in hepatoma H6 treated with brefeldin A for the indicated time periods and (below) quantification. Anti-MBP-ApoL9 antibody was used to detect ApoL9. **D**. qPCR for *Apol9* mRNA levels in hepatoma H6 treated with brefeldin A for the same time periods as in C., normalized to *Gapdh.* Error bars indicate mean ± s.d (n=3). **E-F.** Immunoblotting for ApoL9-V5 in B16F10^L9^ cells treated with EBSS alone, or EBSS+ 400 nM bafilomycin A1, for the indicated time periods. Blots were stripped and reprobed with anti-LC3B antibody to confirm the effect of autophagy inhibition. Blots are quantified and represented as graphs (right). **G**. Immunoblotting for ApoL9-V5 in B16F10^L9^ cells after treatment with 50 mg/mL cycloheximide (CHX) for the indicated time periods and (far right) quantification of the blot. The blot is stained with Ponceau S to show loading.

ApoL9 protein levels in B16F10^L9^ cells were seen to decrease with time during treatment with brefeldin A (**Fig. 3B**). A similar decrease was seen in endogenous ApoL9 in H6 hepatoma cells (**Fig. 3C**). qPCR from H6 hepatoma cells treated with brefeldin A reveals that there is no decrease in ApoL9 mRNA levels during this treatment, indicating that reduced protein levels are not due to a decrease in the transcription of the *Apol9* genes (**Fig. 3D**). Further, amino acid starvation of B16F10^L9^ cells (treatment with EBSS) led to a decrease in ApoL9 levels with time (**Fig. 3E**). To determine if this decrease was due to autophagy, or if ApoL9 was being degraded as a result of the autophagic process, the cells were treated with the autophagy inhibitor bafilomycin A1 in combination with starvation. Bafilomycin A1 was not able to prevent the decrease in ApoL9 protein levels during amino acid starvation (**Fig. 3F**). It is important to note that bafilomycin A1 does not exert its aforementioned effects on the ApoL9 localization pattern during the timescale of the experiment involving starvation with EBSS (6 hours). A similar time-dependent decrease in ApoL9 levels was also noticed when B16F10^L9^ cells were treated with cycloheximide, an inhibitor of protein synthesis (**Fig. 3G**). Therefore, it seems that the decrease in ApoL9 levels during amino acid starvation or treatment with brefeldin A is reflective of normal turnover of ApoL9 in the cell, likely due to decreased protein synthesis in the cell during starvation and ER stress, rather than the result of autophagic degradation. We have previously shown that ApoL9 is primarily degraded by proteasomes, as evidenced by an increase in ApoL9 levels in MG132-treated cells (Arvind and Rangarajan, 2016).

### 4. On treatment with chloroquine, ApoL9 shows a clustered pattern of staining, localises to LAMP1-positive structures and interacts with LC3-II

Like bafilomycin A1, chloroquine (CQ) is also an inhibitor of autophagy. It is a lysosomotropic agent that was thought to accumulate inside lysosomes and reduce their acidity, but it has recently been determined that chloroquine actually inhibits autophagy by preventing autophagosome-lysosome fusion, in addition to having disorganizing effects on the Golgi and endo-lysosomal systems (Mauthe et al., 2018). Treatment of B16F10^L9^ cells with chloroquine resulted in a clustered pattern of ApoL9 in the cytoplasm. Coupling amino acid starvation with CQ treatment (to induce autophagy and simultaneously prevent autophagic degradation) hastened the formation of these clusters and produced a more pronounced pattern for a given time period as compared with treatment with CQ alone (**Fig. 4A**). The ApoL9 in these clusters/vesicles partially colocalized with LAMP1, suggesting that it was interacting with lysosomes or other LAMP1-positive vesicles that are known to be derived from lysosomes (**Fig. 4B**). When a cell line stably expressing V5-tagged LC3B (B16F10^LC3B^) was treated with CQ and EBSS, a similar pattern is observed with LC3B displaying partial but slightly higher degree of co-localization with LAMP1-positive structures, indicating that these are autophagic vesicles, since LC3B is primarily a marker of autophagosomes (**Fig. 4C**). This clearly indicates that ApoL9 is involved in autophagy triggered by amino acid starvation. Treatment with CQ alone does not lead to a build-up of ApoL9 levels in the cell (**Fig. 4D**), nor does CQ prevent the decrease in ApoL9 levels caused by amino acid starvation (**Fig. 4E**). Thus it is unlikely that ApoL9 functions as a cargo receptor that is degraded during selective autophagy.

**Fig.4:**
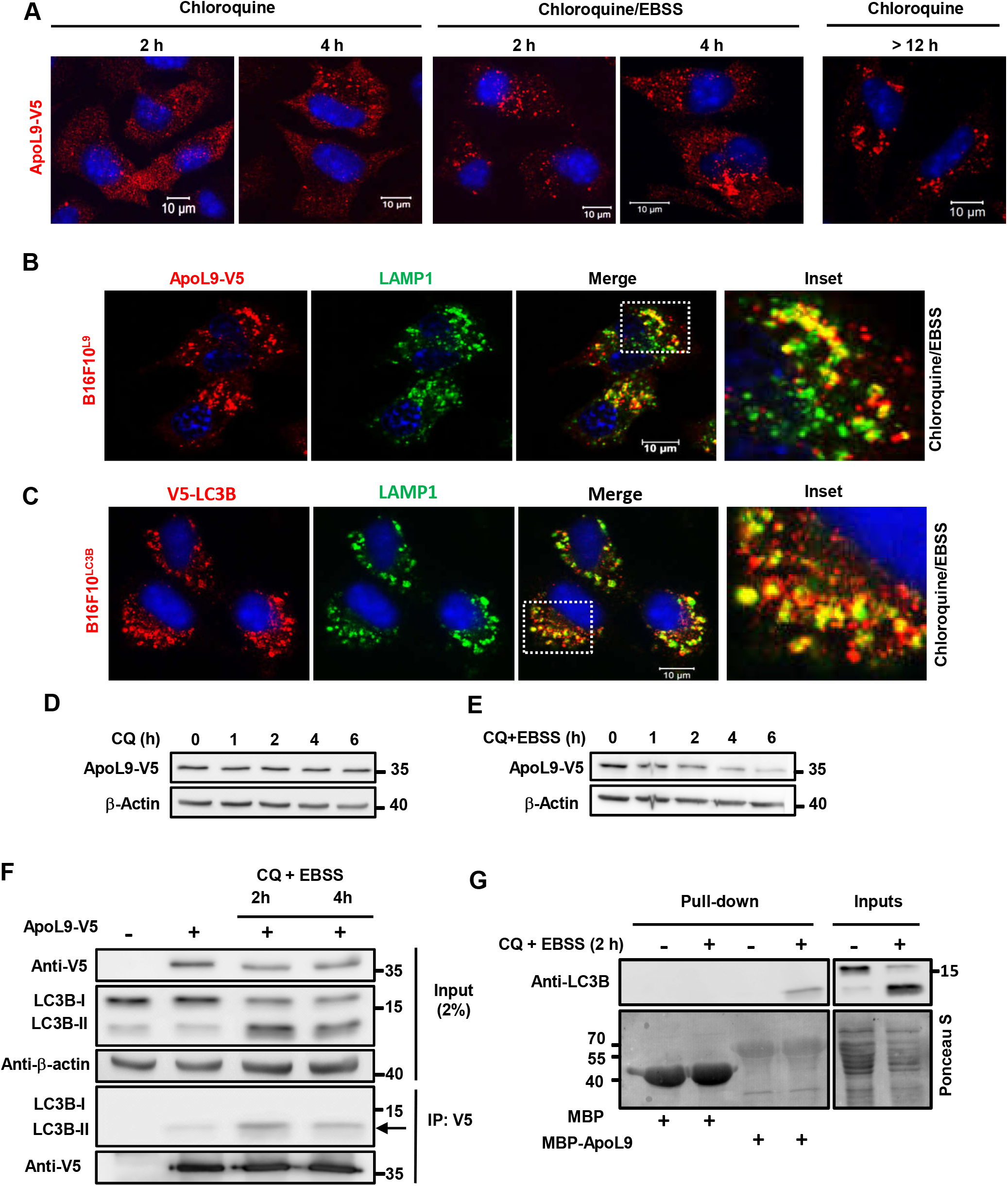
Effect of treatment with chloroquine on ApoL9 in B16F10^L9^ cells. **A.** Subcellular distribution of ApoL9-V5 in B16F10^L9^ cells cultured with 50 mM CQ, or 50 mM CQ+EBSS, for the indicated time periods. Note the heavier aggregation pattern in CQ+EBSS compared with CQ for the same time period. **B, C.** Indirect immunofluorescence for LAMP1 in B16F10^L9^ and B16F10^LC3B^ cells respectively after treatment with CQ+EBSS for 4 h. Note the similar patterns but higher degree of colocalization of LC3B with LAMP1. Boxed insets are magnified on the right. **D, E**. ApoL9-V5 protein levels in B16F10^L9^ cells cultured with CQ alone, or CQ+EBSS, for the indicated time points and (below) quantification. **F.** Co-immunoprecipitation of endogenous LC3B with ApoL9-V5 by anti-V5 agarose from lysates of HEK293T cells transfected with *Apol9-V5* and treated with CQ+EBSS (lanes 3, 4). Lane 1 is cells transfected with control vector, while lane 2 is untreated cells. Note the selective immunoprecipitation of LC3-II in lanes 2-4 (arrow). Anti-LC3B antibody was used. **G.** Pulldown of LC3B from cell lysates of HEK293T cells treated (lanes 2, 4) or not treated (lanes 1, 3) with CQ+EBSS by recombinant MBP/MBP-ApoL9 bound to amylose resin. Blots were stained with Ponceau S to visualize input and recombinant proteins.

Since ApoL9 and LC3B display similar patterns of distribution in amino acid-starved, CQ-treated cells, we decided to examine their interaction during this treatment. HEK293T cells transfected with *Apol9-V5* were subjected to amino acid starvation coupled with chloroquine treatment for different time points to prevent autophagic degradation, thereby causing accumulation of LC3-II. When the cell lysates were incubated with anti-V5 agarose to immunoprecipitate ApoL9-V5, LC3B was co-immunoprecipitated along with ApoL9. Notably, only the LC3-II form was co-immunoprecipitated (**Fig. 4F**), indicating that ApoL9 has a strong affinity for LC3-II. To corroborate this finding, recombinant MBP (maltose binding protein) and MBP-ApoL9 from *E.coli,* bound to amylose resin, were incubated with untreated or chloroquine-EBSS-treated HEK293T lysates. Only LC3-II was pulled down by MBP-ApoL9, confirming that ApoL9 interacts preferentially with that form of LC3 (**Fig. 4G**). This is likely to be a direct result of ApoL9 associating with the PE moiety of LC3-II, since there is otherwise no discernible difference between the two LC3 forms. It is also a clear indication that ApoL9 inside the cell interacts with the lipidated form of LC3 during starvation-induced autophagy, thereby having the potential to influence the process. It must be remembered, from Fig.1A, that ApoL9 has a strong affinity for GABARAP and GABARAPL1 even when these proteins are not in their lipidated form.

### 5a. Mutation of a potential LIR motif _67_FPRL_70_ weakens association of ApoL9 with GABARAP

We have demonstrated that ApoL9 can not only interact with the lipidated form of LC3, but also with nonlipidated LC3 homologues (Fig. 1A).To determine if ApoL9 contains an LIR (LC3-interacting region) motif that is usually required for interaction with LC3/GABARAP family members (Birgisdottir et al., 2013) and has the consensus sequence (W/F/Y)-X-X-(L/I/V), where ‘X’ is any amino acid, we scanned the ApoL9 protein sequence for potential LIRs. Five potential LIRs were identified in the 310 amino acid sequence (**Fig. 5A**). By site-directed mutagenesis, the codons encoding the 1^st^ and 4^th^ (aromatic and hydrophobic) residues of these sequences were mutated to alanines in the ApoL9-V5 construct, generating LIR mutants M1-M5. These mutant proteins were expressed in HEK293T cells and tested for binding to GST-GABARAP. LIR mutants M1, M2, M4, and M5 did not alter binding with GABARAP either when mutated singly (**Fig. 5B**) or when all the four were simultaneously mutated (data not shown). LIR M3, the sequence FPRL (67-70), when mutated to APRA resulted in severely reduced expression of the *ApoL9-V5* construct in cells. When lysate quantities were normalized so as to take equivalent quantities of wild type and mutant protein for the GABARAP pull-down assay, it was seen that mutating this sequence reduced binding to GABARAP by about 50% (average of 3 experiments) (**Fig. 5C**), indicating that the sequence FPRL (67-70) might function as an LIR in ApoL9.

**Fig.5:**
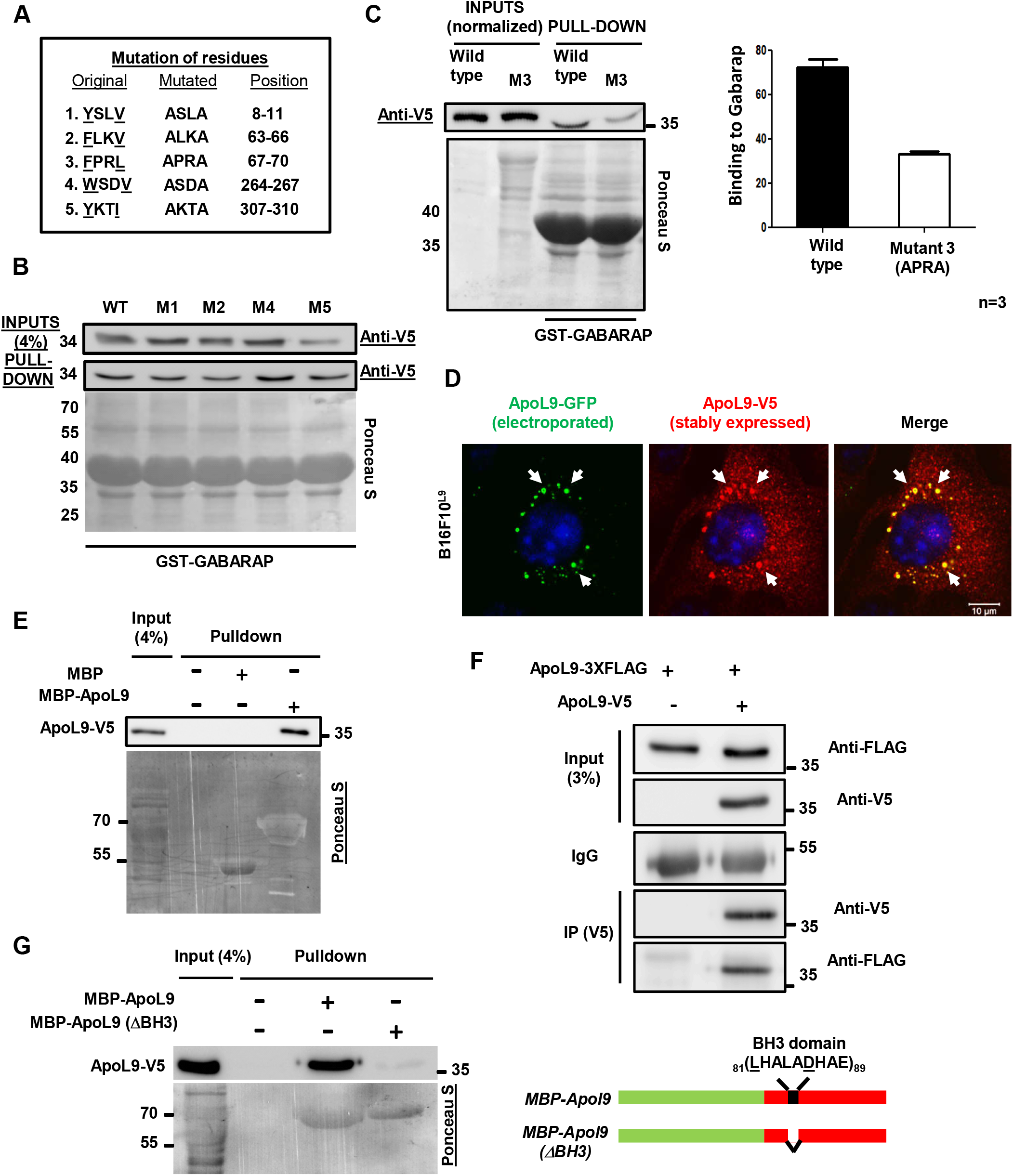
LIR mutants, and oligomerization of ApoL9. **A.** The five potential LIRs, their mutated versions, and their positions in the ApoL9 protein sequence. The mutant forms of the protein were named M1, M2 … M5. **B.** Pull-down of LIR mutants M1, M2, M4 and M5 of ApoL9-V5 (HEK293T) by recombinant GST-GABARAP (from *E.coli),* similar to Fig. 1A. **C**. Comparison of pulldown of WT ApoL9-V5 and LIR mutant M3 by GST-GABARAP, and quantification of binding (mean ± s.d.; n=3) **D**. Altered subcellular distribution pattern of ApoL9-V5 in B16F10^L9^ cells electroporated with ApoL9-GFP (compare with untreated ApoL9-V5 in Fig. 7D/8A). White arrows help visualize similarity in patterns. **E.** Pulldown of ApoL9-V5 from transfected HEK293T cells by recombinant MBP/MBP-ApoL9 bound to amylose resin. Note that only MBP-ApoL9 is able to pull down ApoL9-V5. **F**. Coimmunoprecipitation of ApoL9-3XFLAG with ApoL9-V5 by anti-V5 agarose, from lysates of HEK293T cells transfected with both constructs. Cells transfected with control vector and ApoL9-3XFLAG served as a negative control. **G.** Pull-down of ApoL9-V5 (HEK293T) by recombinant MBP-ApoL9 and MBP-ApoL9 (ΔBH3). A schematic of the BH3 domain deletion is shown on the right.

### 5b. ApoL9 molecules associate with each other

We had reported earlier that the subcellular distribution pattern and morphology of ApoL9-GFP fusions differed greatly from that of ApoL9 with smaller epitope tags, probably due to the larger size of GFP (Arvind and Rangarajan, 2016). It was serendipitously observed that electroporation of *Apol9-GFP* into B16F10^L9^ cells caused the ApoL9-V5 to redistribute in the typical pattern of ApoL9-GFP (**Fig. 5D**), leading us to investigate if both forms of ApoL9 were associating with each other.

When MBP and MBP-ApoL9 bound to amylose resin were incubated with lysate of 293T cells expressing ApoL9-V5, it was determined that ApoL9-V5 associated with MBP-ApoL9 but not with MBP alone (**Fig. 5E**). Further, immunoprecipitation of ApoL9 from lysates of cells transfected with two forms of ApoL9 *(ApoL9-V5* and *ApoL9-3XFLAG)* demonstrated that ApoL9-3XFLAG could be pulled down by ApoL9-V5, indicating that one form of ApoL9 could associate with the other (**Fig. 5F**). The ApoL proteins are BH3-only proteins, as they contain a putative BH3 domain that is characteristic of the Bcl-2 fgamily of proteins (Kreit et al., 2015). Since BH3 domains are known to be involved in homo- and hetero-oligomerization between proteins containing them (Kale et al., 2018), we tested whether deleting the core region of the BH3 domain of ApoL9 (amino acids 81-89) would affect association between ApoL9 molecules. The results indicated that MBP-ApoL9-ΔBH3 was severely impaired in its ability to interact with ApoL9-V5, when compared with wild type MBP-ApoL9. (**Fig. 5G**). This indicated that the BH3 domain is involved in oligomerization of ApoL9 molecules. However, more pull-down experiments *(next section*) reveal that mutations in some other regions also abolish the ability to associate with wild type ApoL9-V5, indicating that the necessary for association/oligomerization might be spread over a wider region of the protein sequence.

### 6. Identification of the PE-binding region of ApoL9

#### 6a. Deletion of amino acids in a putative transmembrane region of ApoL9 abolishes binding to PE

To identify the region of ApoL9 responsible for interaction with PE, we generated a series of mutants carrying deletions in the ApoL9a protein sequence (**Fig. S2.A**). All deletions were made in the MBP-ApoL9 construct cloned in the vector pMAL-c2X. Various criteria were used for making the deletions (**S2.B**). The deletion constructs were expressed as MBP-fusion proteins in *E. coli,* which were purified by amylose affinity chromatography (**Fig. S2.C**). They were then used in protein-lipid overlay assays to examine their ability to bind PE on nitrocellulose membrane, with wild type MBP-ApoL9 serving as a positive control. The results indicated that both the deletions encompassing a putative transmembrane region of the protein (Δ111-123, and Δ124-144; ‘DAS’ prediction server; https://tmdas.bioinfo.se/DAS/index.html) abolished binding to PE (**Fig. 6A**). Results obtained from the two mutants carrying deletions in the BH3 domain and the C-terminal 30 amino acids were inconclusive due to low levels of expression of the recombinant proteins, high levels of impurities during purification and problems with reproducibility.

**Fig.6:**
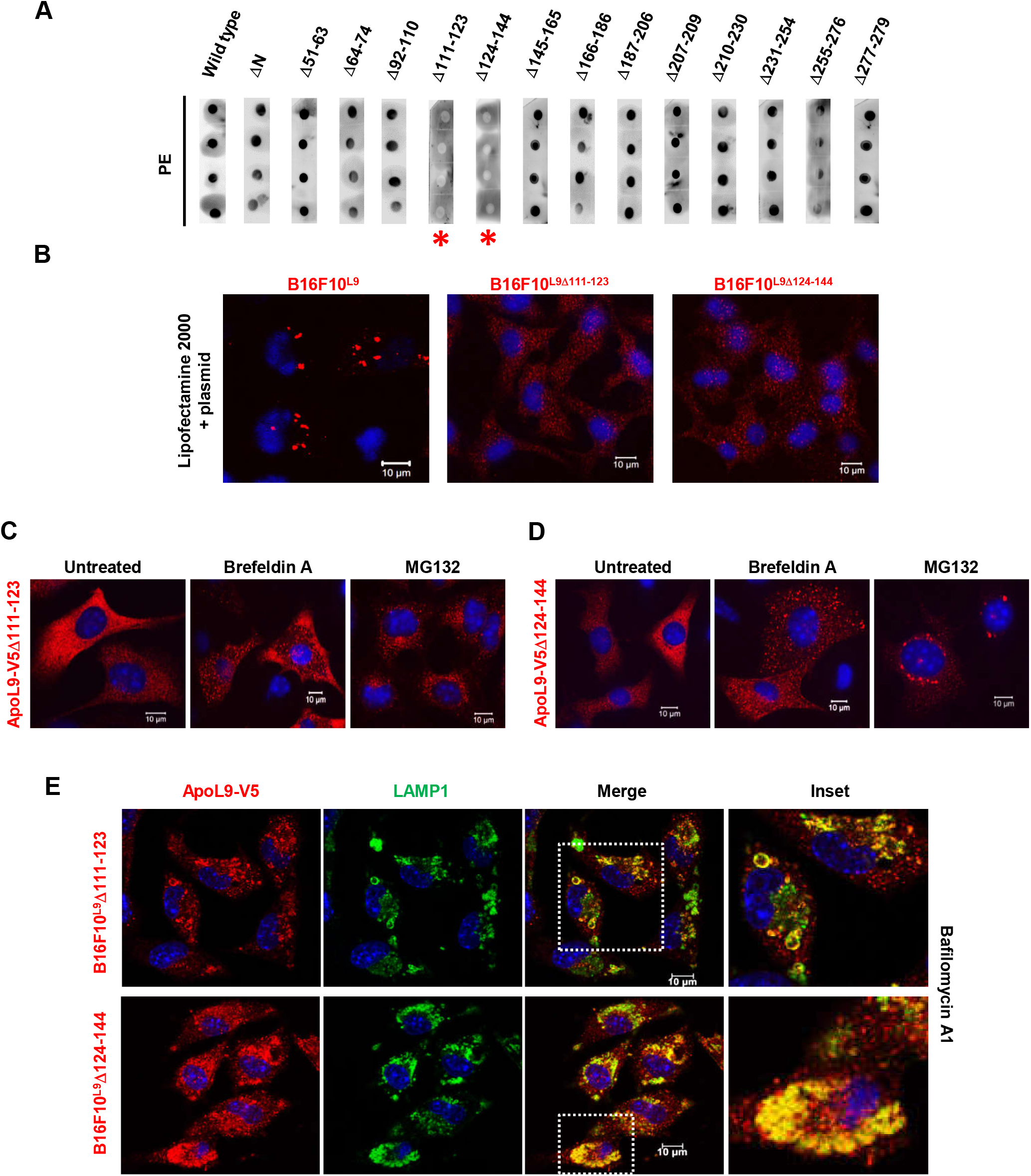
Properties of the region(s) of ApoL9 essential for PE-binding. **A**. Protein-lipid overlay assay for screening PE-binding in MBP-ApoL9 deletion mutants. 1 mg of dipalmitoyl-PE was spotted in quadruplicate on nitrocellulose membrane. Red asterisks indicate mutants that failed to bind PE. **B.** Immunostaining for ApoL9-V5 after lipofection of the insert-less vector *pBApo-EF1 α pur* into cell lines stably expressing wild type ApoL9 or ApoL9 transmembrane deletion mutants for 4 h. Note the characteristic DOPE-induced aggregation only in WT. **C, D**. Subcellular distribution pattern of ApoL9Δ111-123 and ApoL9Δ124-144 electroporated into B16F10 cells and treated with 10 mg/mL brefeldin A for 18 h, or 10 mM MG132 for 6 h. Note the difference in pattern after MG132 treatment. **E.** Indirect immunofluorescence for ApoL9-V5 and LAMP1 in B16F10^L9Δ111-123^ and B16F10^L9Δ124-144^ after treatment with 200 nM bafilomycin A1 for 16 h. Boxed insets are magnified for better viewing.

#### 6b. Non-PE-binding mutants of ApoL9 differ in some of the properties exhibited by wild type ApoL9

We subjected B16F10 cells expressing either ApoL9-V5-Δ111-123 or ApoL9-V5-Δ124-144 to various treatments to see which of the properties displayed by wild type ApoL9 were abolished by deletions in the transmembrane region involvegd in binding PE. We had reported previously that lipofection of any plasmid into B16F10^L9^ cells caused rapid and dramatic aggregation of ApoL9 due to its propensity to interact with dioleoylphosphatidylethanolamine (DOPE) in the transfection reagent (Arvind and Rangarajan, 2016). So, firstly, lipofection of an empty (insert-less) plasmid into cells stably expressing these mutants (B16F10^L9Δ111-123^ or B16F10^L9Δ124-144^) did not result in aggregation of ApoL9, confirming that these mutants were unable to interact with the DOPE in Lipofectamine 2000 (**Fig. 6B**). Secondly, the alterations of ApoL9 distribution in the cells when treated with bafilomycin A1 and brefeldin A, notably the mitochondrial localization of ApoL9 and curved, thread-like structures, were not seen in the mutants (**Fig. 6C, D**). Interestingly, localization to juxtanuclear lysosomes was still observed in bafilomycin A1 treated cells (**Fig. 6E**). Further, the clustered pattern of ApoL9 is not seen when treated with chloroquine-EBSS for shorter time periods, but localization to lysosomes persisted on long-term treatment with chloroquine (**Fig. S3.A**), suggesting that lipid-binding is not linked to lysosomal localization per se, but is still involved in starvation-induced changes in ApoL9 distribution. An interesting observation was that ApoL9-V5-Δ124-144 formed aggresomes on MG132 treatment but ApoL9-V5-Δ111-123 did not (compare **Fig. 6C and 6D**). This was also reflected in the observation that protein levels of ApoL9-V5-Δ111-123 remained more or less unchanged over time on treatment with brefeldin A or cycloheximide, indicating poor turnover, while those of ApoL9-V5-Δ124-144 decreased gradually with time (**Fig. S3.B,C**), suggesting that the region between amino acids 111 and 123 contains residues critical for facilitating proteasomal degradation of ApoL9. Also, interaction with GABARAP remained undisturbed in both the mutants (**Fig. S3.D**), indicating that the transmembrane region had no role in mediating that interaction. On the other hand, both mutant proteins lost the ability to associate with wild type ApoL9 (**Fig. S3.E**), indicating that the intactness of the region is a requisite for oligomerization to happen. A minor note to make is that ApoL9Δ111-123 displays anomalous mobility in SDS-PAGE and migrates slightly slower than wild type ApoL9 despite having a lower molecular mass.

### 7. ApoL9 is a microtubule-associated protein

We know that lipofection of an empty (insert-less) plasmid into B16F10^L9^ cells causes ApoL9-V5 to aggregate in the cytoplasm (Arvind and Rangarajan, 2016; and **Fig. 6B**). But lipofection of an ApoL9-V5-expressing plasmid into B16F10 cells presents a markedly different picture. In addition to a few aggregates, ApoL9 is seen in several long, thin, fibre-like structures (**Fig. 7A**). Further, electroporation of ApoL9-V5 into cell lines such as HepG2 and HEK293T (which express high quantities of protein) also resulted in such structures becoming visible (**Fig. 7B**), ruling out that these structures were another artefact of lipofection of ApoL9. Instead, we think that they are probably formed as a result of high levels of ApoL9 protein expression, since lipofection is a remarkably efficient transfection method that facilitates the expression of very high levels of protein per cell.

**Fig.7:**
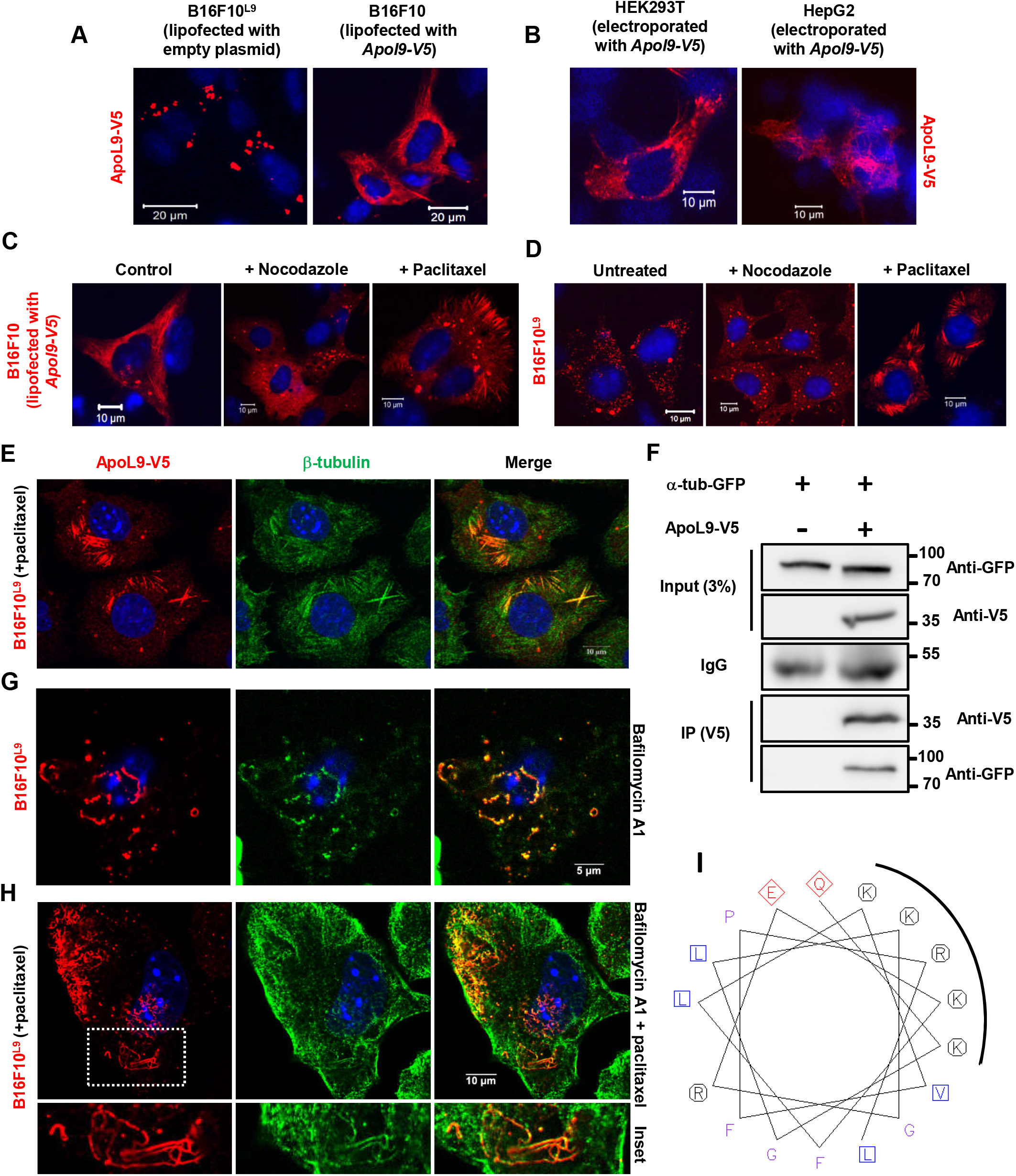
ApoL9 is a microtubule-associated protein. **A.** Difference in distribution pattern of ApoL9 when insert-less plasmid *pBApo-EF1 α pur* is lipofected into B16F10^L9^ cells versus when *Apol9-V5* cloned in *pBApo-EF1 α pur* is lipofected into B16F10 cells. **B.** The *Apol9-V5* construct was electroporated into HEK293T and HepG2 cells. Note the fibre-like structures in the cytoplasm. **C, D.** Effect of compounds that affect microtubule stability on ApoL9 distribution in (C) B16F10 cells lipofected with *Apol9-V5* and in (D) B16F10^L9^ cells. Paclitaxel was used at 10 mM for 4 h and nocodazole at 10 mg/mL for 3 h. **E.** Co-localization of ApoL9-V5 and β-tubulin in B16F10^L9^ cells treated with paclitaxel for 10 h. Anti-b-tubulin antibody was used. **F.** Co-immunoprecipitation of GFP-tagged alpha tubulin with ApoL9-V5 from lysates of HEK293T cells transfected with both constructs. Cells transfected with control vector and the alpha tubulin construct were used as a negative control. **G, H.** Immunoblotting for ApoL9-V5 and β-tubulin in B16F10^L9^ cells treated with bafilomycin (18h) only or with paclitaxel added for the last 8 h. Note the presence of tubulin in curved, thread-like structures. We found tubulin localization in these structures to be sporadic/weak and not seen in all cells. **I.** The region of the helix comprising amino acids 54-70 of mouse ApoL9a, viewed as a helical wheel at http://www.bioinformatics.nl/cgi-bin/emboss/pepwheel. Note the basic residues clustered on one side of the helix.

Since these structures resembled microtubules, we lipofected ApoL9-V5 into B16F10 cells and treated the cells with either nocodazole or paclitaxel (Taxol), a microtubule depolymerizing agent and a microtubule-stabilizing agent respectively. All fibre-like structures disappeared in ApoL9-lipofected cells after a three-hour nocodazole treatment, while their abundance increased (including in the cell periphery) on paclitaxel treatment (**Fig. 7C**), indicating that ApoL9 localizes to microtubules. Further, treatment of the B16F10^L9^ cell line with paclitaxel led to ApoL9-V5 becoming visible on microtubules, which had become stabilized due to the effect of the drug (**Fig. 7D**). Another observation was that treatment of B16F10^L9^ cells with nocodazole greatly increased the number of ALIS-like structures in the cytoplasm (**Fig. 7D**), suggesting that the dynamics of ALIS in the cell are regulated by microtubules and that attachment to microtubules might be involved in trafficking of ApoL9 within the cell. Microtubule association was further confirmed by co-localization of ApoL9 with β-tubulin in paclitaxel-treated cells (**Fig. 7E**). Also, GFP-tagged α-tubulin could be co-immunoprecipitated with ApoL9-V5 from HEK293T cells transfected with both these proteins (**Fig. 7F**), confirming that ApoL9 is a microtubule-associated protein (MAP). Both transmembrane deletion mutants of ApoL9 failed to associate with microtubules, as evident from their immunofluorescence pattern after treatment with paclitaxel, displaying no co-localization with tubulin (**Fig. S4.A**). We have sometimes seen the presence of β-tubulin in the curved, thread-like structures formed during bafilomycin A1 treatment (**Fig. 7G**). This observation has been sporadic and is not readily evident in the majority of cells. But it probably suggests that microtubules are associated with these structures at a certain point in the process of their formation, and might later dissociate from them. It is pertinent to note that LC3B and its orthologues are also microtubule-associated proteins. It has been shown previously that the first 22 amino acids of GABARAP have basic residues clustered on one side of a helix that serves as a tubulin-binding motif (Wang and Olsen, 2000). ApoL9 being a protein that mostly comprises helices in its secondary structure, we examined it for helices enriched in basic residues. The region between amino acids 54 and 70, when visualized as a helical wheel, is part of a helix with at least four basic residues (lysines) clustered on one side (**Fig. 7H**). These lysines are also conserved in the rat orthologue of ApoL9a (**Fig. S4.B**). Further investigations will be needed to reveal if this region indeed binds tubulin and, if yes, to elucidate structural details of the interaction.

### 10. ApoL9 redistributes to stress granules under certain conditions of stress

When B16F10^L9^ cells were subjected to heat stress by incubating them at 43°C for 30 minutes, ApoL9-V5 was seen to redistribute into a large number of granules in the cytoplasm. It was obvious that these granules might be different from ALIS, due to their irregular size and shape, and higher number. In addition to heat stress, we found that other conditions such as oxidative stress induced by sodium arsenite or DTT (but not H_2_O_2_), and ER stress induced by thapsigargin, also caused the redistribution of ApoL9 to such granules (**Fig. 8A**). The identity of the granules was confirmed by co-localization of ApoL9 with the well-known stress granule markers eIF3η (eukaryotic initiation factor 3η) and TIA-1 (T-cell intracytoplasmic antigen 1) (**Fig. 8A, Fig. S4.C**). To confirm that ALIS and stress granules were indeed different, we looked at the localization of SQSTM1 after thapsigargin treatment. Double immunofluorescence showed that SQSTM1 localised to other granules but wasn’t present with ApoL9 in stress granules, confirming that these granules are not ALIS but bona fide stress granules (**Fig. 8C**). Curiously, these two granule types were in close apposition in many locations in the cytoplasm, which could point towards a potential for interaction between their components.

**Fig.8:**
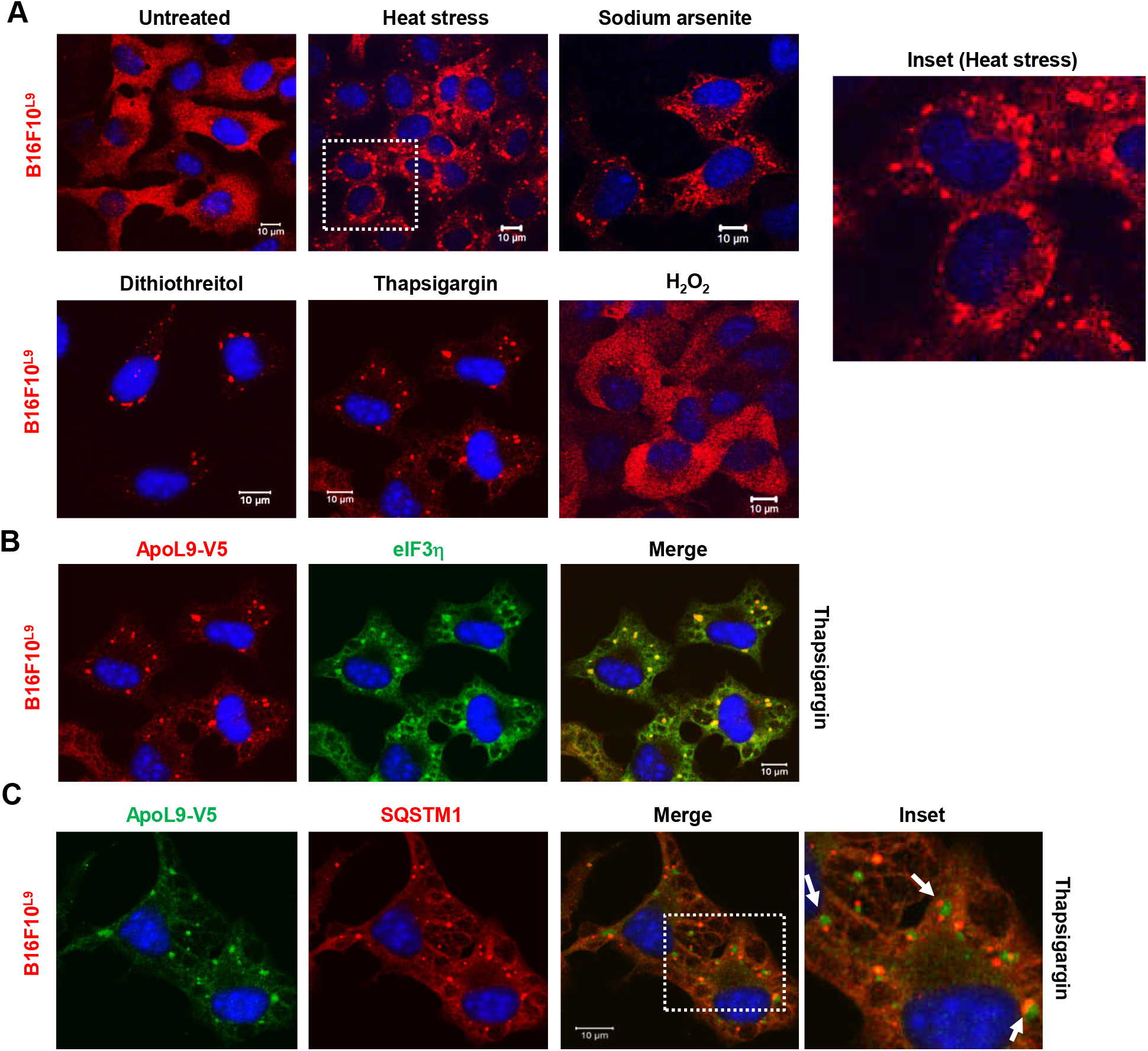
Localization of ApoL9 in stress granules. **A.** Distribution pattern of ApoL9-V5 in B16F10^L9^ cells after treatment with various stress-inducing agents as indicated. Boxed inset (heat stress) is magnified and shown. **B.** Indirect immunofluorescence for ApoL9-V5 and stress granule marker eIF3η in B16F10^L9^ cells treated with 500 nM thapsigargin for 1 h. Anti-eIF3η was used. **C.** Indirect immunofluorescence in thapsigargin-treated B16F10^L9^ cells to distinguish ALIS/sequestosomes (stained with anti-SQSTM1 antibody) from stress granules (stained with anti-V5 antibody). Note the proximity between both types of granules in the magnified inset.

Neither of the transmembrane deletion mutants of ApoL9 (ApoL9^Δ111-123^ or ApoL9^Δ124-144^) redistributed to these granules on subjecting to stress treatments, as evidenced by complete lack of co-localization with eIF3η (**Fig. S.4D**).

## Discussion

Since ApoL9 is a phosphatidylethanolamine-binding protein, we hypothesised that this property is likely to be the biggest determinant of the intracellular functions of the protein. Among the various functions of PE in the cell, its role in autophagy stands out prominently. In this study, we have carried out investigations primarily from this aspect. Each result obtained is discussed in detail in the following paragraphs.

### The possible implications of the ApoL9-LC3/GABARAP interaction

One of the important results in this study is the demonstration of a direct interaction between the lipidated form of LC3 and ApoL9 during starvation-induced autophagy. However, we do not observe ApoL9 concentrated on the numerous punctate autophagosomes that appear in the cell during this process, indicating that it is not an inherent part of the autophagosome. It is likely that ApoL9 transiently interacts with the expanding phagophore/maturing autophagosome. Further, ApoL9 also interacts with the non-lipidated forms of LC3 and its homologues, with highest affinity for GABARAP and GABARAPL1. Though we couldn’t identify an LIR that completely abolished interaction with GABARAP, it is worth noting that the key residues of the LIR motif candidate _67_FPRL_70_, mutation of which reduced binding to GABARAP by >50%, are also conserved in rat ApoL9a. Though the contributions of some other phospholipids (like the various phosphoinosides) in autophagy, membrane remodelling, and mTOR regulation are well catalogued (Dall’Armi et al., 2013), the role of PE in these processes is rather unclear. Shatz et al. state that, in addition to being conjugated with LC3, PE might directly affect lipid bilayer properties of the autophagosome and isolation membrane because it is a cone-shaped lipid and could influence curvature at the leading edge of the forming autophagosome (Shatz et al., 2016). Nath et al. reported a direct correlation between the efficiency of LC3 lipidation and the extent of curvature in lipid bilayers (Nath et al., 2014). Shatz et al. also speculate that proteins with an LIR motif could facilitate docking of phagophore membrane precursors by binding to LC3 on the formed phagophore or the incoming membrane during phagophore formation. In this context, ApoL9 is unique in that it is capable of interacting with non-lipidated LC3/GABARAP, lipidated LC3/GABARAP, with PE alone, as well as with other ApoL9 molecules, allowing a multidimensional range of possibilities as to how it might influence events at the membrane during autophagosome formation. It would also be worth investigating the effect of ApoL9 itself on membrane curvature, since it is associated with highly curved structures during treatment with bafilomycin A1. Hsu and Shi state in their review of mitochondrial phospholipids in health and diseases that lipids, unlike proteins and nucleic acids, have been stubbornly difficult to visualize in cells, and “the next leap in the understanding of autophagic membrane dynamics requires direct visualization of phospholipids in living cells to capture the transient autophagosomes” (Hsu and Shi, 2017).

### Could ApoL9 influence mitophagy?

The fact that the PE-binding ability of ApoL9 is abolished by the deletion of a putative transmembrane domain is directly indicative of the likelihood of the protein binding to PE embedded in lipid bilayers/membranes. Localization to swollen mitochondria during treatment with brefeldin A and bafilomycin A1 is also abolished in these deletion mutants ApoL9Δ111-123 and ApoL9Δ124-144. Since the bulk of the PE in the cell is synthesised in the mitochondria by decarboxylation of phosphatidylserine (Borkenhagen et al, 1961; Shiao et al., 1995), it is not surprising to find ApoL9 in mitochondria (though it is not detected there in steady state conditions). The outer mitochondrial membrane (OMM) is known to have a plethora of pro-and anti-apoptotic proteins of the Bcl-2 family that regulate mitochondrial integrity and physiology (Harris and Thompson 2000). We have observed ApoL9 forming complete rings around mitochondria, and incomplete ‘crescents’ around some (Fig. 2A, 2D, and S4.E), suggesting the possibility that ApoL9 molecules might gradually colonize the mitochondrial membrane. This is reflected in a few images where the parts of the membrane densely stained for TSPO are weakly stained for ApoL9, and vice versa (**Fig. S4.E**). It is common knowledge that autophagosomes are of different sizes depending on the cargo that is targeted for degradation. For quite some time, it has been known that the mitochondria are one of the various sources that contribute membrane to form autophagosomes by membrane addition (Hailey et al., 2010; Cook et al., 2014). Cook et al. observed through electron microscopy that mitochondrial membranes were sometimes contiguous with the membranes of the forming autophagosome. Hailey et al. speculate about how the curvature-inducing property of PE at the mitochondrial membrane, together with the hemifusion-inducing property of LC3, could potentially be important to generate autophagosomes from mitochondria. They also note that during autophagy, a significant quantity of membrane moves from autophagosomes to lysosomes through fusion, and that PE comprises a large fraction of this. To the best of our knowledge, this study reports the first instance of a PE-binding protein that has been observed localizing to mitochondria synchronously with proteins of the autophagic machinery, and co-localizing with them. Another connection worth noting is that ApoL9 has been reported to interact with PHB1 and PHB2 (Kreit et al., 2015), two proteins of the inner mitochondrial membrane (IMM) that have been implicated in controlling IMM organization by acting as protein and lipid scaffolds (Osman et al., 2009). Further, PHB2 has been identified to function as a receptor for mitophagy on the IMM by bridging LC3 and depolarized mitochondria whose outer membranes have ruptured (Wei et al., 2017). How ApoL9 could fit in this complex scheme of events is a topic for future study.

It has been reported (Maeda and Fadeel, 2014) that intact mitochondria could be expelled from cells undergoing TNF-α-induced necroptosis and that this appeared to happen before the disruption of the plasma membrane, suggesting an active mechanism of release. Several other reports have also suggested that molecules derived from mitochondria and other organelles can act as DAMPs (damage-associated molecular patterns) that act as signals for other cells in the immune system (Krysko et al., 2011; Zhang et al., 2010). In our experiments, we have observed ApoL9-containing ring-like structures being released from cells as early as 12 hours post bafilomycin A1 treatment, in which period the cell membranes are unlikely to have suffered much damage from the actions of the compound. Sun et al. (Sun et al., 2014) had reported that secretion of ApoL9a from cells can induce epithelial proliferation, but commented that the mechanism of secretion was unclear. It would be pertinent to investigate if ApoL9 can be secreted as part of the mitochondrial membrane in other cell types. That the ApoLs are TNF-α-inducible genes points towards such a possibility.

### Oligomerization of ApoL9

Several cellular phenomena are regulated by proteins that oligomerize. This property might confer additional advantages to proteins in certain conditions, such as improved stability, the ability to concentrate molecules spatially, better interaction with partner proteins, and to assemble complexes and scaffolds (Marianayagam et al., 2004). We can envisage a scenario where ApoL9, by interacting with other ApoL9 molecules, can group together lipid molecules (PE) or proteins like lipidated LC3 and concentrate them on discrete regions of the lipid bilayer on membranes of autophagic vesicles, or organelles like lysosomes. Interorganellar contacts involving oligomerization of ApoL9 could also be possible, for example between autophagosome-lysosome, or between mitochondria. Several of the images of mitochondria show dense spots containing ApoL9 on the membrane, and the mitochondria are also sometimes clustered together in apparent close contact with each other, with the points of contact between mitochondria showing dense ApoL9 staining (Fig. 2B, brefeldin). Though we do not know the significance of ApoL9 localizing to stress granules at the moment, several proteins that are found in stress granules, e.g. TIA-1, TIAR, G3BP, etc. undergo oligomerization with the help of some or other specific type of domain contained within them (Tourriere et al., 2003; Gilks et al., 2004; Hua and Zhou, 2004).

The mutants ApoL9Δ111-123 and ApoL9Δ124-144 do not exhibit several of the properties displayed by the wild type protein. Of the many properties of wild type ApoL9, association with mitochondria and formation of insoluble curved structures seem to be the most likely properties linked with the ability to bind PE, due to involvement of membrane/membrane fractions. At this stage, it is difficult to comment on whether stress granule localization or microtubule binding depends on interaction with lipid. However, oligomerization, interaction with GABARAP, and localization to aggresomes are not directly linked to lipid-binding.

### In what way could a lipid-binding MAP influence autophagy?

The ability of ApoL9 to associate with microtubules might influence several cellular functions of the protein, because it can potentially ferry lipid, or other, molecules to various parts of the cell while attached to microtubules. Like several other MAPs (Itoh and Hotani, 1994), ApoL9 seems to stabilize microtubules by binding to them, since mere overexpression of ApoL9 is sufficient for stabilized microtubules to become visible in the cytoplasm. ApoL9 seems to be directly and strongly associated with microtubules unlike proteins like LC3B that associate indirectly by virtue of their association with other MAPs, since treatment of the B16F10^LC3B^ cell line with paclitaxel does not result in LC3B becoming visible on stabilized microtubules in the cytoplasm (**Fig. S4.F**). We could co-immunoprecipitate α-tubulin with ApoL9, but since tubulin forms heterodimers further investigations are needed to determine which tubulin types ApoL9 associates with and what the structural details of the interaction are. It is interesting to observe that the helical region comprising amino acids 54-70 that is a potential tubulin-binding motif lies exactly adjacent to FPRL, the potential LIR motif of ApoL9 which when mutated reduced association with GABARAP by >50%. GABARAP itself is a tubulin-binding protein, and it is not inconceivable that ApoL9 might simultaneously associate with microtubules and GABARAP under certain conditions. Autophagosomes and lysosomes are known to specifically travel along microtubules that undergo specific types of post-translational modifications (Xie et al., 2010; Mohan et al., 2018); therefore it might be of interest to investigate if ApoL9 localization is restricted to some or other subset of microtubules.

The identity of the highly curved, thread-like structures that accumulate in the cytoplasm after prolonged treatment with bafilomycin A1 is unclear, as they did not co-localize with any protein or organelle marker we used. Since their increase in the cell correlates with ApoL9 fractionating into a Triton X-100-insoluble fraction of the lysate, it is possible that they comprise only ApoL9 and membranous material. ApoL9 localizes to lysosomes on treatment with bafilomycin A1 or CQ, and this property is not abolished in the non-PE-binding mutants suggesting that PE-binding is not necessary for lysosomal localization. The C-terminus of ApoL9 contains the sequence YKTI, which corresponds to a known consensus for lysosomal membrane targeting (YXXΦ) (Guarnieri et al., 1993). ApoL9 seems to be targeted to lysosomes only under certain conditions, because in normal conditions it doesn’t seem to accumulate in these organelles. This points to the possibility of ApoL9 being involved in the later stages of autophagy, when autophagosomes mature and autolysosomes come into play. The thread-like structures do not accumulate in cells expressing the non-PE-binding mutants; it is likely that these structures are formed as a result of the prolonged downstream block in the autophagic process due to the action of bafilomycin A1. It is widely accepted now that the terminal step of autophagy is a process called autophagic lysosome reformation, in which a clathrin-based budding mechanism facilitates the growth and elongation of tubules from autolysosomes along microtubules, followed by scission of proto-lysosomes (Rong et al., 2012; Du et al., 2016). However, it is still very unclear how different phospholipids regulate this mechanism, and how lysosomal membrane and luminal proteins are sorted in this process. In our study, as mentioned before, we have noticed a negative correlation between the presence of ApoL9 in juxtanuclear lysosomes and in curved thread-like structures. We have also occasionally observed microtubule proteins co-localizing with these structures. Therefore, we speculate that ApoL9, being a MAP and an oligomerizing PE-binding protein, might regulate the formation of these or other similar tubules from the lysosomes, thus potentially recycling membrane components for autophagy. It is interesting to note that these ApoL9-positive curved, thread-like structures are seen only in bafilomycin A1-treated cells but not in CQ-treated cells, and that the positioning of lysosomes in the former case tends to be predominantly juxtanuclear. This may reflect the differences in the mechanism of autophagy inhibition by these two compounds.

Taken together, the data presented in this study demonstrates that ApoL9 is a rather dynamic protein that is found in several distinct compartments in the cell and interacts with various cellular molecules. It is reasonable to speculate that it is involved in lipid transport in the cell and that its role in autophagy is likely to be that of an adaptor protein. Since it is restricted to two species of rodents and its expression levels are tissue-specific, it is unlikely to be an essential protein for autophagy and we propose that it might confer some very selective advantage to rodent-specific metabolic activities. One of the most significant differences between bigger and smaller animals is the high metabolic rate per unit weight in the latter (Klieber, 1947). Assuming that ApoL9 has a small but not insignificant contribution in augmenting a process such as protein homoeostasis or autophagy, and considering its high levels in liver and brain tissue where autophagy plays critically important roles in maintaining organ homoeostasis, it is possible to envisage a scenario where expression of such a gene could confer a definite advantage to the fitness of a fast-metabolising species such as mouse. Therefore we also feel that it would be appropriate if further investigations regarding ApoL9 be performed in an *in vivo* context where its functions might be understood even better, like in hepatic or neuronal tissues and in knockout mice.

## MATERIALS AND METHODS

### Cell lines and reagents

B16F10 mouse melanoma was obtained from the tissue culture facility, Dabur Research Foundation, Ghaziabad, India. Mouse hepatoma H6 has been described (Thompson et al., 1981) and referenced (Nandi et al., 1996) previously. HEK293T (ATCC^®^ CRL 3216™) and HepG2 (ATCC^®^ HB-8065™) were gifts from Prof. Ganesh Nagaraju and Prof. Anjali Karande, Indian Institute of Science, Bangalore, India. All cell lines were cultured in high glucose Dulbecco’s Modified Eagle’s Medium (DMEM) containing 10% foetal bovine serum (FBS) and an antibiotic mixture comprising 10,000 units/mL penicillin, 10,000 μg/mL streptomycin, and 25 μg/mL amphotericin B. The cell lines were maintained in an incubator at 37°C with 5% CO_2_ and water provided for humidification. All cell lines were routinely inspected for the absence of any potential contamination, including mycoplasma.

DMEM, FBS (E.U.-approved, South America origin), Earle’s Balanced Salt Solution (EBSS), Opti-MEM reduced serum medium, Antibiotic-Antimycotic solution (100X), Pierce BCA protein assay kit, and PageRuler prestained protein ladder were purchased from Thermo Fisher Scientific, Waltham, Massachusetts, USA. Trypsin (from porcine pancreas), puromycin dihydrochloride, Lipofectamine 2000, TRI reagent for RNA isolation, chloroquine, thapsigargin, paclitaxel, nocodazole, MG132, DTT and 1,2-Dipalmitoyl-sn-glycero-3-phosphoethanolamine were purchased from Sigma-Aldrich, St. Louis, Missouri, USA. H_2_0_2_ and sodium arsenite were purchased from Merck, Darmstadt, Germany.

Bafilomycin A1 and brefeldin A were purchased from Cell Signaling Technology, Inc. (CST), Danvers, Massachusetts, USA. Glutathione resin was purchased from G-Biosciences, St. Louis, Missouri. Bovine albumin, fraction V, fatty acid-free was purchased from MP Biomedicals Ltd., Navi Mumbai, Maharashtra, India. Amylose resin and all restriction endonucleases were procured from New England Biolabs (NEB), Ipswich, Massachusetts, USA. All oligonucleotides used in PCR and cloning were procured from Sigma-Aldrich. XT-20 high fidelity polymerase was used in all cloning protocols and purchased from GeNei Pvt. Ltd., Bangalore, India. D (+) Maltose monohydrate was purchased from HiMedia Laboratories Pvt. Ltd., Mumbai, India. Calf thymus DNA was procured from Amersham plc, Buckinghamshire, United Kingdom. cOmplete™ protease inhibitor cocktail tablets, EDTA-free, were purchased from Roche, Basel, Switzerland.

### Antibodies

The following antibodies were used in this study: anti-V5 (mouse monoclonal, R960-25, Thermo Fisher Scientific), anti-FLAG (mouse monoclonal, F1804, clone M2, Sigma-Aldrich), anti-LC3B (#2775, rabbit polyclonal, CST), anti-β-actin (A3854, rabbit polyclonal, HRP-conjugate, Sigma-Aldrich), anti-V5 agarose affinity gel (A7345, monoclonal clone V5-10, Sigma-Aldrich), anti-eIF3η (sc-16377 (N20), goat polyclonal, Santa Cruz Biotechnology, Inc., Dallas, Texas, USA), anti-GFP (sc-9996, mouse monoclonal clone B2, Santa Cruz), anti-SQSTM1 (Ab56416, mouse monoclonal, Abcam, Cambridge, United Kingdom), anti-V5 (V8137, rabbit polyclonal, Sigma-Aldrich), anti-LAMP1 (clone 1D4B, rat monoclonal, Developmental Studies Hybridoma Bank, University of Iowa, Iowa, USA). Rabbit polyclonal anti-MBP-ApoL9 antibody has been described and validated earlier in (Arvind and Rangarajan, 2016). Also it should be noted that, in the bulk of experiments in the paper, ApoL9-V5 was detected in immunostaining and immunoblotting by anti-V5 antibody, unless specifically mentioned otherwise.

### Plasmids and constructs

The *Apol9a* gene was amplified as described earlier (Arvind and Rangarajan, 2016) from cDNA prepared from total RNA of B16F10 cells treated with type I interferon. ORFs for other genes were amplified from cDNA prepared from various mouse tissues/cell lines according to level of expression. The *Apol9-V5* and *Apol9-3XFLAG* constructs were made by cloning the cDNAs into the XbaI and HindIII sites in the vector *pBApo-EFlαpur* (Takara Bio Inc., Kusatsu, Shiga Prefecture, Japan). The V5 sequence was included in the reverse primer, while the 3XFLAG sequence was amplified from the vector *p3XFLAG-CMV-10* (Sigma-Aldrich). The *MBP-ApoL9* construct was made by cloning *Apol9a* amplified from the *Apol9-V5* construct and cloning into the SalI and HindIII sites in the vector *pMAL-c2X* (NEB). MBP and MBP-ApoL9 were transformed into *E.coli BL21 (DE3)* competent cells and purified according to the manufacturer’s instructions. The inserts for *Lc3b* (BamHI/SalI), *Lc3a* (EcoRI/SalI), *Gabarap* (BamHI/SalI), *Gabarapl1* (EcoRI/SalI), and *Gabarapl2* (EcoRI/SalI) were amplified from cDNA derived from B16F10 melanoma cells and cloned in the vector *pGEX-4T-1* (GE Healthcare, Chicago, Illinois, USA) into the sites indicated in parentheses. *Lc3b* was also cloned with an N-terminal V5 tag into the BamHI and HindII sites of *pBApo-EF1α pur.* To create GFP-fusion constructs *Lc3b* (EcoRI/BamHI) and *Gabarap* (HindIII/BamHI) were also cloned into the vector *pEGFP-C1* (Clontech Laboratories, Mountain View, California, USA). N- and C-terminal GFP fusions of ApoL9 were created by cloning *Apol9a* into *pEGFP-C1* (HindIII/KpnI) and *pEGFP-N1* (HindIII/ApaI) respectively. The plasmid expressing GFP-tagged alpha tubulin was a gift from Prof. Vaishnavi Ananthanarayanan, Centre for BioSystems Science and Engineering, IISc, Bangalore, India. Translocator protein *(Tspo)* was cloned into the HindIII and BamHI sites of pEGFP-N1. *Tia-1* was cloned into the EcoRI and BamHI sites of *p3XFLAG-CMV-10.*

### LIR mutants and deletion constructs

LIR mutants of *Apol9-V5* were created by using complementary primers containing the desired mutations in the region of the potential LIR, using wild type *Apol9-V5* to create amplicons with an overlapping sequence, followed by overlap extension PCR to make the final mutant construct, and cloning into the HindIII and XbaI sites of *pBApo-EF1α pur.*

Similarly, deletions in the *MBP-Apol9* sequence were made by using complementary primers containing a deletion in the region of the desired deletion, followed by overlap extension PCR. The constructs were cloned into the SalI and HindIII sites in the vector *pMAL-c2X.* The WT *MBP-Apol9* construct was used as template for the initial rounds of PCR. To create *Apol9-V5 (Δ111-123)* and *Apol9-V5 (Δ124-144),* the inserts were amplified from the respective *MBP-Apol9* constructs and cloned into the XbaI and HindIII sites of *pBApo-EF1α pur.* All clones were sequenced to confirm for the desired mutations or deletions.

The various deletions are described and shown schematically (**Fig. S2.A,B**). The N-terminal deletion was 54 amino acids, spanning the full length of the helices in that region. The C-terminal deletion was 30 amino acids in length which corresponded to a predicted coiled coil sequence in that region. The BH3 deletion mutant encompassed amino acids 81-89, the core BH3 domain consensus sequence. Based on predictions from software that predict transmembrane sequences in proteins (Cserzo et al., 1997), two deletions were made that encompass potential transmembrane regions on the ApoL9a sequence – amino acids 111-123, and 124-144. The other deletions were made randomly in the remaining regions of the sequence, with no deletion exceeding an average length of 20-25 amino acids. Two exceptions were made based on exact stretches of 3 amino acids being present in mouse PEBP1 and human PEBP4, respectively, both being proteins of the phosphatidylethanolamine-binding protein family.

### Creation of cell lines stably expressing protein

The B16F10 cell line stably expressing ApoL9 (B16F10^L9^) has been described previously (Arvind and Rangarajan, 2016). Briefly, the *ApoL9-V5* construct in *pBApo-EF1α pur* was linearized using ScaI and transfected into semi-confluent B16F10 cells using Lipofectamine 2000. 48 h later, the cells were subcultured into six-well plates and transformants were selected with 2 μg/mL puromycin. Ten days later, single colonies were picked using sterile PYREX^®^ cloning cylinders (6 mm x 8 mm, Corning Inc., Corning, New York, USA) and expanded into cultures in 6-well plates. The colonies were screened for ApoL9 expression by indirect immunofluorescence using anti-V5 antibody, and clones with an optimum level of expression were further expanded and frozen. B16F10^LC3B^, B16F10^L9Δ111-123^ and B16F10^L9Δ124-144^ were made by a similar procedure, using the *V5-LC3B, ApoL9-V5Δ111-123* and *ApoL9-V5Δ124-144* constructs.

### Transfection procedures

Lipofectamine 2000 was used according to the manufacturer’s instructions. Electroporation was done using Bio-Rad Gene Pulser™ electroporation cuvettes (0.4 cm gap; blue) and the Gene Pulser Xcell™ electroporation system. Briefly, confluent B16F10 cells in a T75 culture flask were trypsinized and centrifuged at 350 *g.* The cells were resuspended homogeneously in 2.5 mL Opti-MEM reduced serum medium. 400 μL of cell suspension was mixed with 5-10 μg of plasmid DNA and ~20-30 μg of sterile, sonicated calf thymus DNA, added to the cuvette and pulsed using the exponential decay protocol at 240V and 950μF capacitance. The electroporated cell suspension was added to 1.2 mL Opti-MEM in 35 mm culture dishes with or without glass cover slips. The medium was replaced with fresh medium 4-6 h post electroporation, and the cells were processes for downstream experiments 16-24 h post transfection.

Calcium phosphate transfection was performed as described in (Sambrook and Russell, 2005) with minor modifications. This technique was primarily used for HEK293T cells. Briefly, the growth medium of cells (70% confluent) was replaced with 4 mL Opti-MEM just prior to transfection. For transfection of a 60 mm dish, a cocktail was made that contained 8 μg plasmid DNA, ~30 μg calf thymus DNA, and 22 μL of 2M CaCl_2_ made up to 180 μL with water. To this, 180 μL of 2X Hepes-buffered salt solution was added drop-wise with intermittent slow aeration by pipetting. The mixture was incubated for 20 minutes and then added to the cells gently. The transfection medium was replaced with fresh medium exactly 4 h post transfection, and the cells were processed for downstream experiments ~24 hours after transfection.

### Cell lysis and protein estimation

All cells were lysed in situ, i.e. by adding lysis buffer to the cell monolayer on the culture dishes and incubating in a rocker at 4 °C. For making total cell lysates, RIPA buffer (50 mM Tris-HCl pH 7.4, 150 mM NaCl, 0.1% SDS, 0.5% sodium deoxycholate, 1% NP-40, 1 mM EDTA and complete protease inhibitor cocktail) was used. The lysate was briefly sonicated to shear viscous DNA and centrifuged at 10,000 rpm at 4 °C in a tabletop microcentrifuge. The supernatant was used for western blotting. For making Triton X-100 soluble and insoluble fractions of cell lysates, a lysis buffer comprising 50 mM Tris-HCl pH 7.4, 150 mM NaCl, 1 mM EDTA and 0.1% Triton X-100 with protease inhibitor cocktails was used. The pellet was resuspended in an appropriate volume of Laemmli buffer, after centrifugation to clarify the lysate at 10,000 rpm at 4 °C on a microcentrifuge. The supernatant was used for western blotting, pull-down, IP, or other experiments. All protein estimations were performed using the Pierce BCA protein assay kit (Thermo Fisher Scientific).

### SDS-PAGE and western blotting

SDS-PAGE gels were cast using a pre-made solution of acrylamide:bisacrylamide :: 29:1. Cell lysates were loaded on gels, in most cases at 30 μg total protein per lane. The resolved proteins on the gels were transferred to PVDF membranes (FluoroTrans^®^ W, 0.2 μm pore size, Pall Corporation, New York, USA) by semi-dry transfer at constant current. Recipes for separating and stacking gels, gel running buffer and transfer buffer were standard recipes from (Green and Sambrook, 2014). Membranes were blocked in 5% fat-free milk prepared in TBST (Tris-buffered saline with Tween 20; 50 mM Tris-HCl pH 8.0, 150 mM NaCl, 0.1% Tween 20). Primary antibodies were added according to the concentration provided by the manufacturer (usually a dilution of 1: 1000 – 1: 5000), and secondary HRP-conjugated antibodies were used at a dilution of 1: 10000. The blots were developed using Immobilon^®^ chemiluminescent HRP substrate (EMD Millipore, Merck), on an ImageQuant LAS 4000 platform (GE Healthcare). Quantitation of western blots was done using the software ImageJ, and graphs were plotted using GraphPad Prism, version 6 (GraphPad Software, California, USA). Blots were stripped using a buffer comprising 62.5 mM Tris-HCl pH 6.8, 2% SDS, and 100 mM 2-mercaptoethanol, and re-probed for loading controls and other proteins.

### Protein pull-down and co-immunoprecipitation

For GST/MBP pull-down assays, recombinant GST-fusion/MBP-fusion constructs were transformed into BL21-DE3 *E. coli*, and glycerol stocks of colonies were made at a final concentration of 15% glycerol. The cultures were inoculated into LB medium and induced with 0.5 mM IPTG for 3.5 h at 37°C. After centrifugation, the pellets were lysed with a lysis buffer (20 mM Tris-HCl, 200 mM NaCl, 1 mM EDTA, pH 7.4, with added 1 mM PMSF). The solution was sonicated to lyse the cells and clarified in a cold centrifuge at 10,000 *g.*

30-50 μL of affinity resin (amylose/glutathione agarose) was washed with the bacterial lysis buffer twice, and the required quantity of bacterial lysate was added to it. The lysate was incubated on an end-to-end rotator for binding for 30 min to 1 h, followed by three washes with wash buffer (lysis buffer with a tiny amount of Triton X-100 for smooth movement of liquid in tube). Next, cell lysate expressing/overexpressing the protein of interest was incubated with the beads for 2 h to overnight, followed by three washes with wash buffer (mammalian cell lysis buffer). The beads were then boiled in Laemmli buffer, centrifuged, and the supernatant was used for SDS-PAGE, followed by western blotting. A fraction of the cell lysates used for IP was loaded as input. GST/MBP protein was detected by staining the western blot membrane with a solution of 0.1% Ponceau S in 5% acetic acid immediately after transfer.

### Indirect immunofluorescence

Cells grown on acid-washed coverslips were fixed with 3.7% formaldehyde, washed twice with PBS, and permeabilized with 0.1% Triton X-100 in PBS. After blocking with 5% BSA in PBS for 30 min., they were incubated with primary antibodies in blocking solution for 2 hours. All antibodies were used at 1:200 dilution, except anti-LAMP1 (1:50). After 3 washes with PBS containing 0.05% Triton X-100, they were incubated with secondary fluorophore-conjugated antibodies (Alexa Fluor 555/488; Thermo Fisher Scientific/CST; 1:200 dilution) in blocking solution for 2 hours. After 3 more washes, they were stained with 1μg/mL DAPI or Hoechst 33342, washed once, and mounted on a drop of ProLong^®^ series antifade reagents (Thermo Fisher Scientific).

Imaging was done on either of these platforms: Zeiss LSM 510, Zeiss LSM 880 with Airyscan, or Leica TCS SP8 confocal microscopes. Intensity, brightness and colour contrast of images were balanced, and scale bars were drawn, using either of the following software depending on which microscope was used for imaging: LSM Image Browser or ZEN 2.3 Black (Carl Zeiss Microimaging), or Leica Application Suite X ver. 3.4. Cropping, arranging and labelling of images were done using a combination of Adobe Photoshop, Microsoft PowerPoint, and GNU Image Manipulation Program (GIMP).

### Treatment of cells in culture

Bafilomycin A1 and brefeldin A were dissolved in DMSO to make stock solutions of 100 μM and 10 mg/mL respectively. DMSO alone was used as a vehicle for treatment of cells in control dishes. For all cell treatments, the addition of DMSO to cells never exceeded 0.1%. Chloroquine was dissolved in water to make a 10 mM stock solution. Heat stress was applied by keeping cells in a CO_2_ incubator set at 43°C for 30 min. Sodium arsenite was added to cells at 0.5 mM for 30 min. DTT was added to the cells at 2 mM, from a freshly prepared stock solution in water. Thapsigargin was used at 500 nM for 1 h, from a 1.5 mM stock solution in methanol. Bafilomycin A1 was used at 200 nM for appropriate time periods, unless mentioned otherwise. Brefeldin A was used at 10 μg/mL. Chloroquine was used at 50 μM. EBSS treatment of cells was done by washing the cells with EBSS twice after removing the normal culture medium, and culturing in fresh EBSS for the stipulated time periods. Paclitaxel was dissolved in DMSO to make a 10 mM stock and used to treat cells at 10 μM. Nocodazole was dissolved in DMSO to make a 10 mg/mL stock and used to treat cells at 10 μg/mL. MG132 was dissolved in DMSO to make a 10 mM stock and used to treat cells at 10 μM for 6 hours. Cycloheximide was dissolved in water to make a 10 mg/mL stock and used at a concentration of 50 μg/mL in the chase assay.

### Protein-Lipid overlay assay

MBP-ApoL9 and its deletion mutants were purified by binding cell lysates of IPTG-induced *E.coli* cultures to amylose resin, eluted with 20mM maltose and quantified using a NanoDrop (Thermo Fisher Scientific). They were electrophoresed on an SDS-PAGE to check for protein integrity. 1,2-Dipalmitoyl-sn-glycero-3-phosphoethanolamine was dissolved in chloroform to 1 mg/mL and warmed to dissolve fully. 1 μl of the solution was spotted on a nitrocellulose membrane (BioTrace™ NT, Pall Corporation) in quadruplicate and allowed to dry for 30 minutes. The membrane was blocked in TBST containing 3% fatty acid-free BSA overnight, followed by incubation with the respective purified MBP-fusion protein (2 μg/mL) for 2 h. After 3 washes in TBST, the membranes were incubated with primary antibody (anti MBP-ApoL9 rabbit polyclonal) at 1:3000 dilution and secondary antibody (goat anti-rabbit IgG-HRP) at 1:10000 in blocking solution and, as in the procedure for western blotting, washed, coated with luminol substrate and viewed in a chemiluminescence imager.

## Acknowledgements

The staff of the Confocal/Bioimaging Facility, Division of Biological Sciences, Indian Institute of Science, are gratefully acknowledged for their assistance.

