## Supplementary Figures for "Apolipoprotein L9 interacts with LC3/GABARAP and is a microtubule-associated protein with a widespread subcellular distribution"

#### Supplementary Figure 1

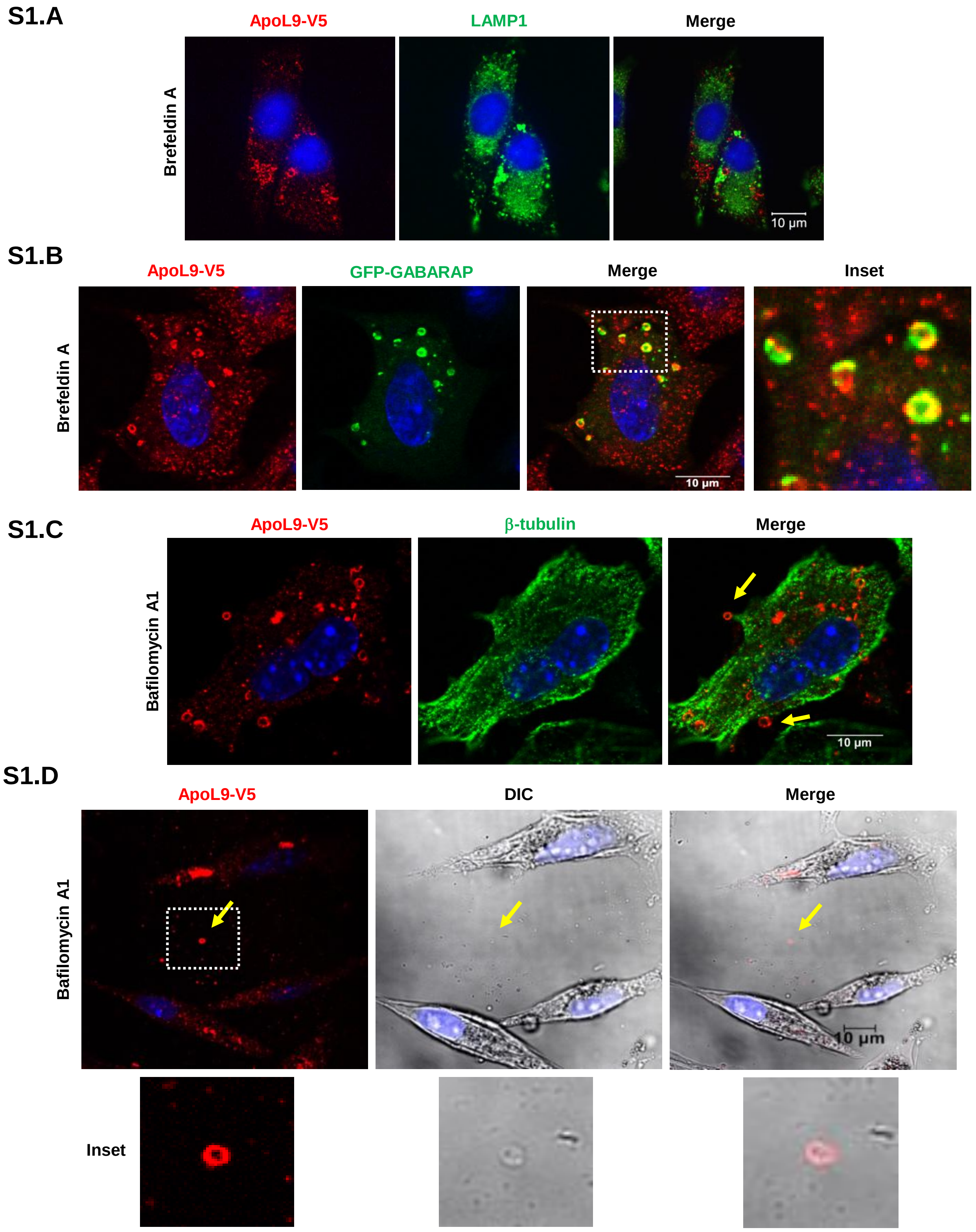

**Fig.S1:**

**A.** Indirect immunofluorescence for ApoL9-V5 and LAMP1 in B16F10$^{L9}$ cells treated with 10 mg/mL brefeldin A for 16 h. Note the lack of co-localization of ApoL9 with LAMP1. **B.** Indirect immunofluorescence for ApoL9-V5 and GFP-GABARAP in B16F10$^{L9}$ cells treated with 10 mg/mL brefeldin A for 16 h. Note the presence of GABARAP on/near all structures (mitochondria) positive for ApoL9. Boxed inset is magnified for better viewing. **C.** Ring-like structures positive for ApoL9-V5 exiting a cell treated with bafilomycin A1 for 16 h (yellow arrows). Staining for tubulin (green) helps visualize the cell boundary. **D.** Released ring-like structure in the extracellular region (yellow arrow) and magnified boxed inset. Note the vesicle-like appearance of the same in the DIC image.

### Supplementary Figure 2

## S2.A

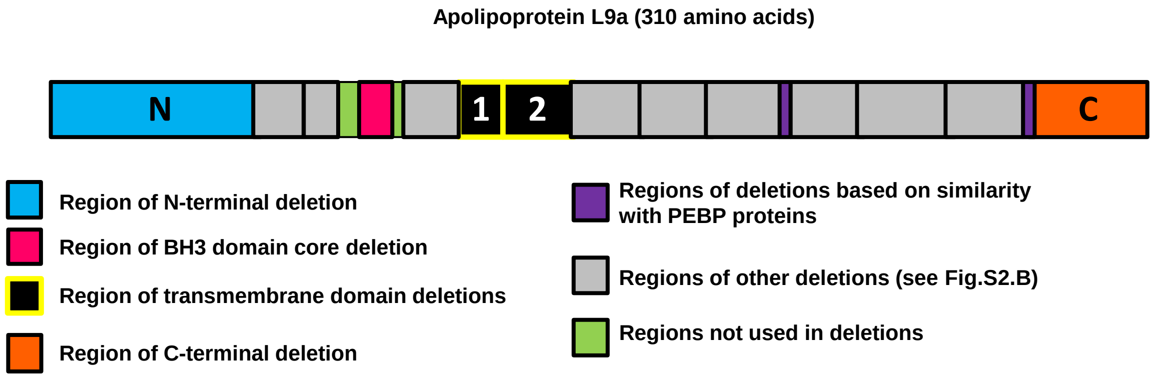


## S2.B

| Deletion | Residues | Reasoning behind deletion | Deletion | Residues | Reasoning behind deletion |
|---|---|---|---|---|---|
| ΔN | 1-54 | N-terminal helices | Δ166-186 | 166-186 | Random deletion |
| Δ51-63 | 51-63 | Random deletion | Δ187-206 | 187-206 | Random deletion |
| Δ64-74 | 64-74 | Random deletion | Δ207-209 | 207-209 | Conserved identity with human PEBP4 |
| ΔBH3 | 81-89 | Core BH3 consensus | Δ210-230 | 210-230 | Random deletion |
| Δ92-110 | 92-110 | Random deletion | Δ231-254 | 231-254 | Random deletion |
| Δ111-123 (TM1) | 111-123 | Part of predicted TM domain | Δ255-276 | 255-276 | Random deletion |
| Δ124-144 (TM2) | 124-144 | Part of predicted TM domain | Δ277-279 | 277-279 | Conserved identity with mouse PEBP1 |
| Δ145-165 | 145-165 | Random deletion | ΔC | 281-310 | Predicted coiled coil domain |

## S2.C

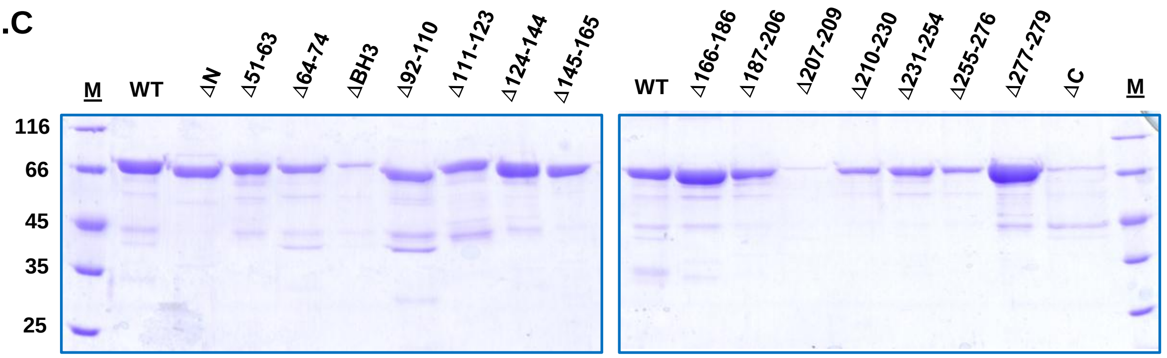


**Fig.S2:**

**A.** Schematic of deletions constructed in the ApoL9 coding sequence to screen for PE-binding. **B.** Table detailing the deletions made in the ApoL9 coding sequence and the reasoning behind the various deletions. **C.** MBP-ApoL9 deletion mutant proteins resolved by SDS-PAGE and stained with Coomassie Brilliant Blue

### Supplementary Figure 3

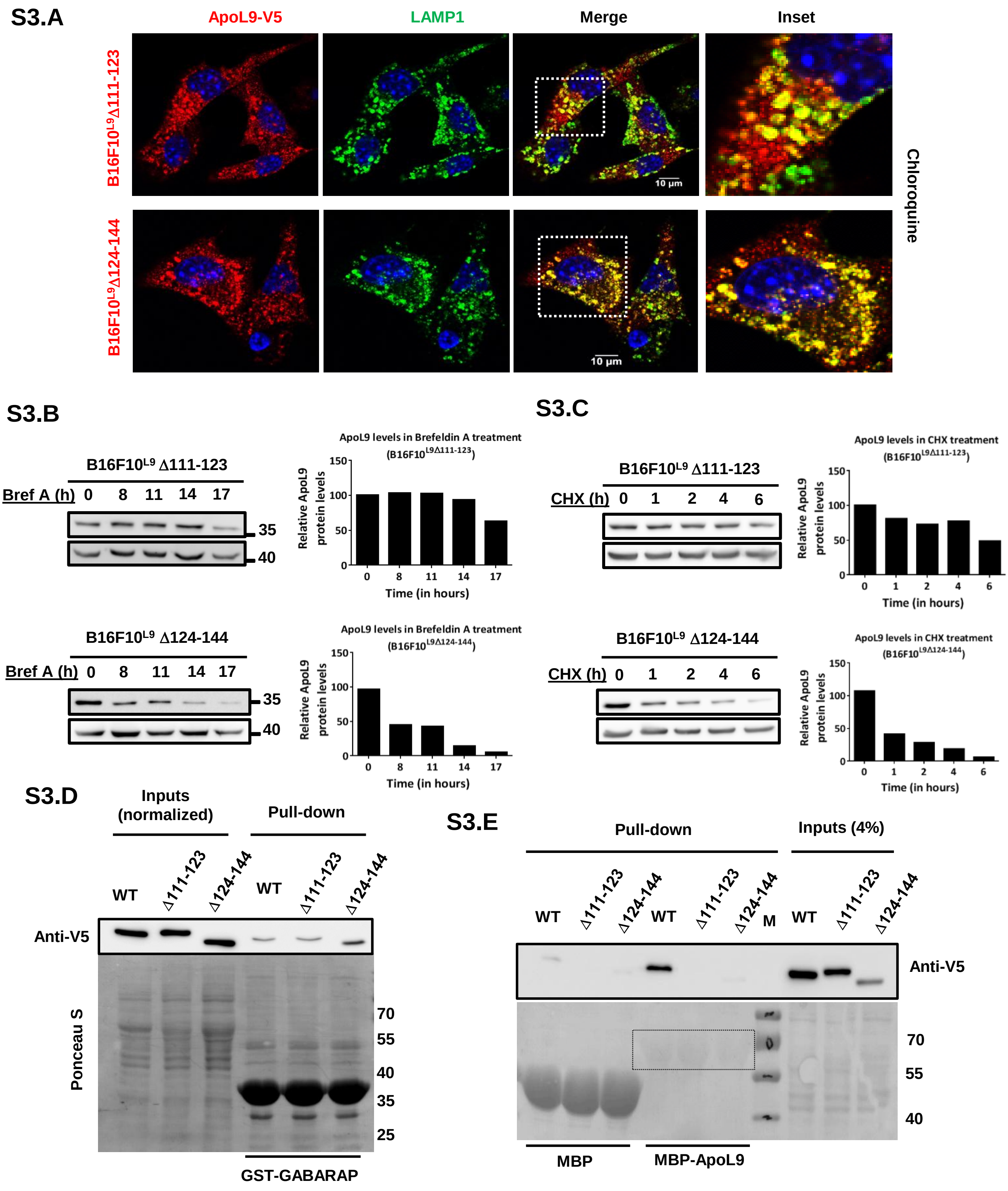

**Fig.S3:**

**A.** Indirect immunofluorescence for ApoL9-V5 and LAMP1 in B16F10$^{L9D111-123}$ and B16F10$^{L9D124-144}$ cells treated with chloroquine for 12 hours. Boxed insets are magnified and shown. Anti-V5 and anti-LAMP1 antibodies were used. **B.** ApoL9-V5 levels in B16F10$^{L9D111-123}$ and B16F10$^{L9D124-144}$ cells treated with brefeldin A for the indicated time periods and (right) quantification of the same. Note the distinct decrease in ApoL9 levels only in B16F10$^{L9D124-144}$. **C.** ApoL9-V5 levels in B16F10$^{L9D111-123}$ and B16F10$^{L9D124-144}$ cells treated with 50 mg/mL cycloheximide for the indicated time periods and (right) quantification of the same. Anti-V5 antibody was used for immunoblotting. **D.** Pull-down of ApoL9$^{D111-123}$-V5 and ApoL9$^{D124-144}$-V5 (from HEK293T lysates) by recombinant GST-GABARAP, similar to Fig. 1A. WT ApoL9-V5 binding to GST-GABARAP serves as a positive control. Levels of input were normalized such that equal quantities of mutant proteins were used in the assay. Recombinant GST-GABARAP is stained with Ponceau S. **E.** Pull-down of ApoL9$^{D111-123}$-V5 and ApoL9$^{D124-144}$-V5 (from HEK293T lysates) by recombinant MBP and MBP-ApoL9 bound to amylose resin. WT ApoL9-V5 binding to MBP-ApoL9 serves as a positive control. MBP and MBp-ApoL9 are shown stained by Ponceau S. Loading of MBP-ApoL9 is lesser than that of MBP, hence it is shown boxed.

### Supplementary Figure 4

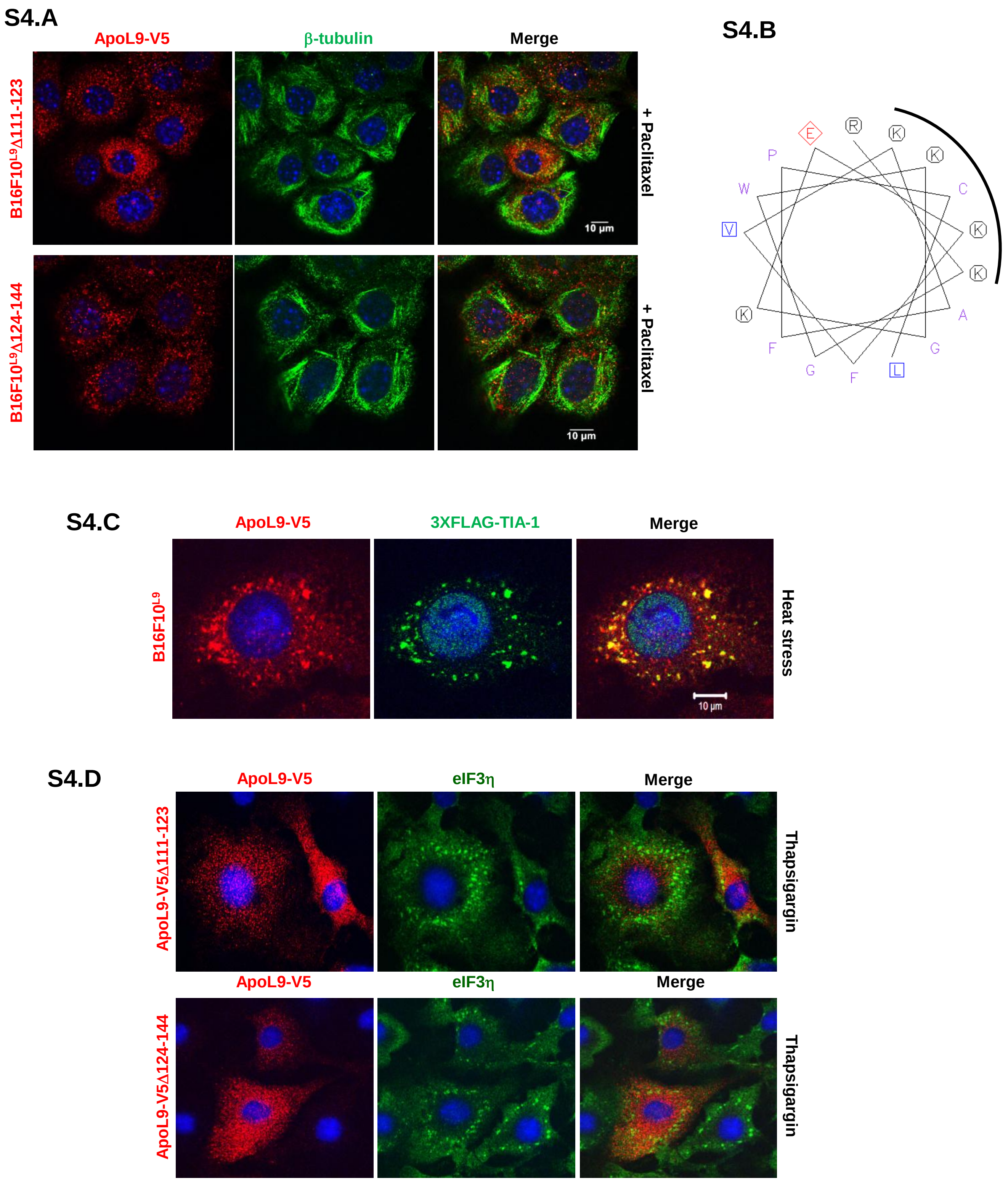

S4.E

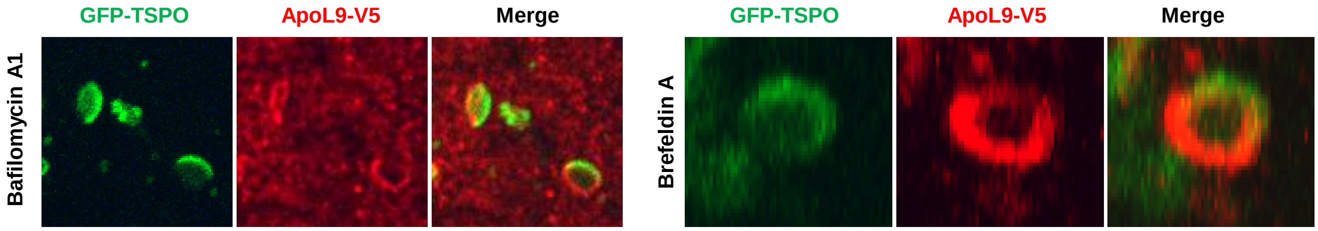


S4.F

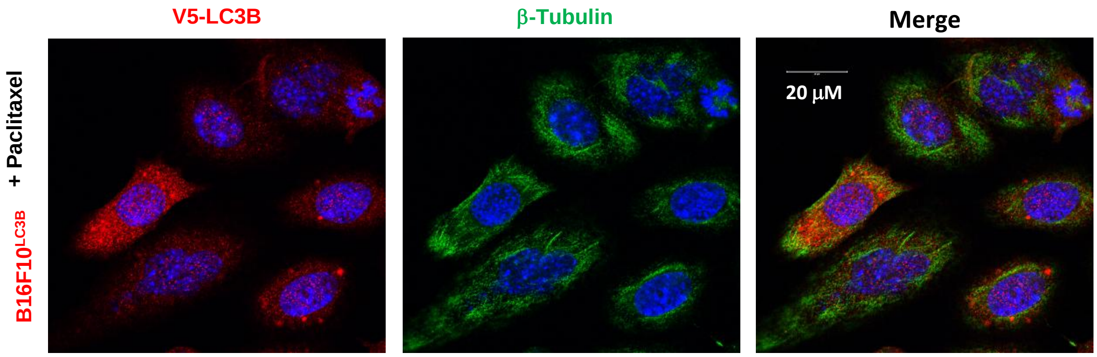


**Fig.S4:**

**A.** Indirect immunofluorescence for ApoL9-V5 and b-tubulin in B16F10$^{L9D111-123}$ and B16F10$^{L9D124-144}$ cells treated with paclitaxel for 10 h. Note the lack of association of ApoL9 with microtubules. **B**. The region of the helix comprising amino acids 54-70 of rat ApoL9a, viewed as a helical wheel at http://www.bioinformatics.nl/cgi-bin/emboss/pepwheel . Note the conserved lysines by comparing with mouse ApoL9a in Fig. 7H. **C.** Indirect immunofluorescence for ApoL9-V5 and stress granule marker 3XFLAG-TIA-1 in B16F10$^{L9}$ cells treated at 43˚C for 30 min. The *3XFLAG-TIA-1* construct was electroporated into B16F10$^{L9}$ cells. Anti-V5 and anti-FLAG antibodies were used. **D.** Indirect immunofluorescence for ApoL9$^{D111-123}$-V5 and ApoL9$^{D124-144}$-V5 and eIF3h in B16F10 cells electroporated with the respective ApoL9 deletion mutants and treated with 500 nM thapsigargin for 1 h. Neither of the mutants is present in stress granules. **E.** Magnified images of selected ApoL9-positive mitochondria in B16F10$^{L9}$ cells treated with bafilomycin A1/brefeldin A. Note how parts of the mitochondria densely stained for TSPO-GFP are weakly stained for ApoL9-V5, and vice versa. **F.** Indirect immunofluorescence for V5-LC3B and b-tubulin in B16F10$^{LC3B}$ cells treated with paclitaxel for 10 h. Note that LC3B is not evidently visible on microtubules (compare with Fig. 7E).
